# Selection can favor a recombination landscape that limits polygenic adaptation

**DOI:** 10.1101/2024.08.22.609166

**Authors:** Tom Parée, Luke Noble, Denis Roze, Henrique Teotónio

## Abstract

Meiotic crossover positions are uneven along eukaryotic chromosomes, giving rise to heterogeneous recombination rate landscapes. Genetic modifiers of local and genome-wide crossover positions have been described, but the selective pressures acting on them and their potential effect on adaptation in already-recombining populations remain unclear. We performed experimental evolution using a mutant that modifies the position of crossovers along chromosomes in the nematode *Caenorhabditis elegans*, without any detectable direct fitness effect. Our results show that when the recombination landscape is fixed, adaptation is facilitated by the modifier allele that, on average, increases recombination rates in genomic regions containing heritable fitness variation. However, in polymorphic populations containing both the wild-type and mutant modifier alleles, the allele that facilitates adaptation tends to decrease in frequency. This is likely because the allele that reduces recombination between selected loci at the genome-wide scale increases recombination in its chromosomal vicinity, and may thus benefit from local associations it establishes with beneficial genotype combinations. These results demonstrate that indirect selection acting on a recombination modifier mainly depends on its local effect, which may be decoupled from its consequences on genome-wide polygenic adaptation.

## Introduction

Crossovers are characteristic of sexual eukaryotes, ensuring that parental genotypes recombine while preventing homolog chromosome missegregation during meiosis [1]. Although most eukaryotes only have one or two crossovers per bivalent at each meiosis, their positioning along the chromosomes is not uniform, resulting in recombination rate “landscapes” that may vary between species, populations, individuals, or even between sexes [2–9]. Recombination landscapes are known to correlate with patterns of genomic organization, such as gene density, transposon activity, and chromatin structure [10–13], or patterns of genetic diversity, such as levels of heterozygosity and genetic differentiation within and among species [14–16]. Yet, the selective forces affecting the evolution of recombination landscapes have remained elusive.

Recombination modifiers can be under direct selection when they cause aneuploidies through chromosomal non-disjunction or compromise DNA repair, which usually reduces fertility and/or increases embryonic lethality [17–19]. More interestingly, in the presence of negative linkage disequilibrium (LD) between selected loci, genetic modifiers increasing recombination rates are expected to facilitate adaptation by expanding the pool of standing genetic variation available to natural selection and disrupting the associations between beneficial and deleterious alleles [20–25]. Negative LD may result from the deterministic effects of negative epistatic interactions among selected loci [26–28], or from the stochastic interference effect among selected loci in finite populations known as the Hill-Robertson effect [29–32]. In agreement with these theoretical expectations, several experiments indicate higher adaptive rates and lower effects of linked selection and selective interference in sexual (and recombining) populations than in asexual ones [33–36]. The benefits of increased recombination rates to adaptation in already-recombining populations are nonetheless expected to be more modest [23, 31, 37, 38], and have been challenging to demonstrate experimentally [39, 40]. A modifier increasing average recombination rates across the whole genome should have more effect on adaptation when the genetic architecture of fitness is highly polygenic [37, 41], which often seems to be the case in natural populations [42, 43]. However, theoretical results have shown that indirect selection on recombination modifiers should be mostly driven by the associations they establish with beneficial genotypes in their local genomic neighborhood [24, 44], and be less affected by their effect on recombination between more distant loci.

We here study if and how a genetic modifier of crossover position across the genome impacts adaptation, and the resulting indirect selection acting at the modifier locus, using the *rec-1* gene of the nematode *Caenorhabditis elegans* as an experimental evolution model. At each *C. elegans* meiosis, there is exactly one crossover per pair of homologous chromosomes for the six holocentric chromosomes, with a reduced probability of crossover occurrence in the large central chromosomal regions relative to flanking regions [45]. This recombination landscape is shared with related nematodes [46–48], separated from *C. elegans* for up to tens of millions of years, and correlates with the genome organization and the genetic diversity found in natural populations [49–52]. Loss-of-function mutants of the *rec-1* gene redistribute crossover positions to central chromosomal regions, smoothing the recombination landscape without changing crossover number [53–56].

## Results and Discussion

### Experimental framework

Three sets of evolution experiments were performed to study the evolution and adaptive consequences of the *rec-1* recombination modifier (Figure S1). In the first set of experiments, direct selection on alternative *rec-1* alleles was estimated to test for potential effects of the recombination modifier on fertility and embryo survival [17–19]. Direct selection was measured in populations without genetic diversity across the genome, except at the *rec-1* locus, so meiotic crossovers occurred without effective recombination. A second set of experiments examined the effect of the recombination landscape on adaptation to a new environment, in genetically diverse populations fixed for alternative *rec-1* alleles. The third set of experiments estimated indirect selection on segregating *rec-1* alleles in genetically diverse populations, in which meiotic crossovers lead to effective recombination of genetic variation.

The populations used in these experiments were derived from a population previously domesticated to standard laboratory conditions, referred to here as the domestication environment. The domesticated population was initially derived from the hybridization of 16 founder strains and cultured at high population size (census size *N* = 10^4^, effective size *N_e_*= 10^3^) and predominant outcrossing between hermaphrodites and males for 140 generations [57–59]. It contains abundant standing genetic variation with genomic heritability for hermaphrodite self-fertility (a fitness-proxy) of 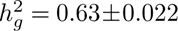 SD (see Methods). At the same time, linkage disequilibrium between founder alleles remains high only at relatively short genetic distance (0.5 cM on average) [60, 61]. The three evolution experiments presented here were conducted in the domestication environment and a novel environment characterized by a 9-fold increase in the NaCl concentration of the growth media. The novel salt environment decreases hermaphrodite fertility and male mating ability (Figure S2), presumably by retarding larval development to maturity [59].

### No direct selection at *rec-1*

Two pairs of wild-type and mutant lines were obtained by editing loss-of-function *rec-1* mutants in two isogenic lines derived from the domesticated population [56](Figure S1, Table S1). To quantify direct selection at the *rec-1* locus, experimental evolution was performed with the *rec-1* wild-type and mutant alleles competing in otherwise isogenic backgrounds. The initial sex ratio was also varied to account for potential *rec-1* allele fitness differences between male and hermaphrodite meioses, and the possibility for frequency-dependent selection was further controlled. In total, 143 replicate populations were cultured for five generations at *N* = 10^3^ (Table S3).

During the experiments, changes in frequencies of *rec-1* alleles should be explained by their fitness differences (measured as a direct selection coefficient of the *rec-1* mutant allele, *s_d_*; see Methods). We find that *rec-1* alleles have similar fitness as *s_d_* is not different from 0 (Figure 1; Generalized Linear Model, *s_d_* = 1.47 *×* 10*^−^*^3^, bootstrap p-value = 0.758; see Methods), independently of the environment and initial sex ratio (environment p=0.352, sex-ratio p=0.159; Figure S3). Although its effect is significant, the isogenic background has little influence on the relative fitness of the *rec-1* alleles (p-value=4.6 *×* 10*^−^*^3^; Figure S3). Hence, there is little opportunity for direct selection at the *rec-1* locus under the environmental and demographic conditions employed. These findings confirm previous studies showing that *rec-1* loss-of-function mutants do not affect embryonic survival and fertility [53, 55, 56].

**Fig. 1.**
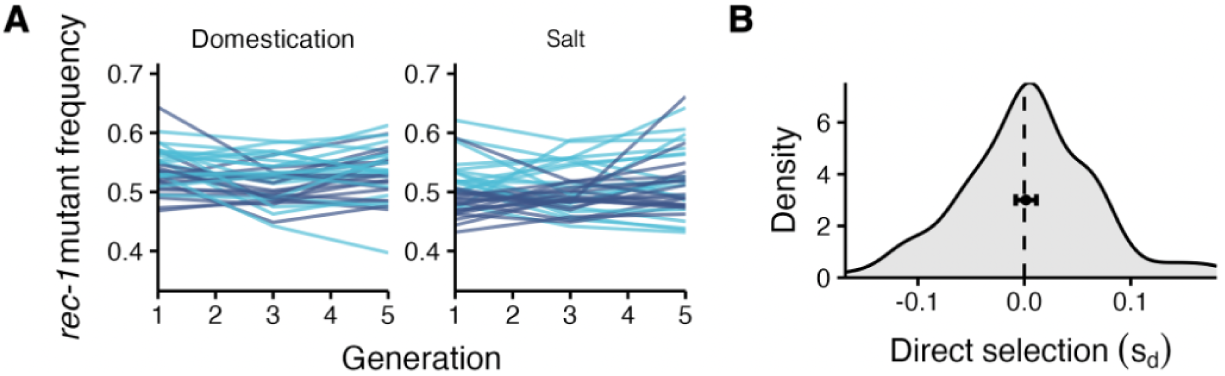
Direct selection at the *rec-1* locus. **A.** *rec-1* mutant allele frequency during five generations of competition between isogenic lines differing only in *rec-1* allele identity (n=143 replicate populations at *N* = 10^3^; 66 replicates at different initial allele frequencies not shown). Without genetic diversity, these experiments test for the fitness effects of compromised DNA repair or chromosome non-disjunction. Competitions were conducted in two different environments (domestication or salt), varying the genotype background (EEV1401 vs EEV1403, light blue; EEV1402 vs EEV1404, dark blue; see Table S1 for strain designation and definition), and the sex ratio (high or low male frequency; not shown). During the experiments, *rec-1* mutant allele frequency was measured every other generation using a qPCR protocol (Figure S13; Methods). **B.** Kernel density distribution (smoothed histogram) of direct selection coefficients of the *rec-1* mutant allele (*s_d_*) in each replicate, as estimated by a binomial GLM regression of allele frequency on generations of competition (see Methods). The dashed line indicates 0. The black dot shows the *s_d_* GLM generation estimate and thus the average relative fitness of the *rec-1* allele over all replicate populations and conditions. Error bar is the 95% credible intervals (CI) estimated by 10^4^ bootstraps with resampling of replicates. The effect of each treatment is shown in Figure S3.

### Recombination landscapes correlate with fitness heritability

While the *rec-1* mutant allele does not have an obvious direct fitness effect, it could impact adaptation by changing recombination rates of genomic regions containing fitness variation. Genetically diverse sister populations, differing in *rec-1* alleles (homozygous wild-type or mutant), were obtained by mass-introgression of one of the *rec-1* loss-of-function mutants into the domesticated population (Figure S1, Table S1, see Methods). Re-sequencing samples of individual pools from these sister populations indicated that 90.8% of the single nucleotide variants (SNV) from the domesticated population were retained and that SNV frequencies are similar between the sister populations (Figure S4A, Table S2, *r*^2^ = 0.92). The exception is a region surrounding *rec-1* on the left of chromosome I where, as expected, the introgressed allele containing the *rec-1* mutant is more frequent in the *rec-1* mutant sister population (Figure S4B). However, this difference has no consequence for the results and conclusions presented here (see below).

The genomic regions where the mutant decreases the probability of crossovers and thus reduces recombination rates (Δ*r <* 0; Figure 2A) have higher genetic diversity (Figure 2B). This is because SNV diversity is higher in genomic regions that recombine more than the rest of the genome under the wild-type *rec-1* allele (in particular, near the extremities of chromosomes). The effect of the *rec-1* mutant on recombination rate landscapes of the sister populations is thus inversely correlated with standing genetic variation (Figure 2C). By reducing recombination in the most diverse genomic regions, the *rec-1* loss-of-function mutant increases proportion on SNV that rarely recombine (Figure 2D), and we predict that the *rec-1* mutant should increase the average genome-wide linkage disequilibrium between SNVs (Figure 2E). In addition, and using a large collection of isogenic lines derived from the domesticated population [60, 61], we find that less SNV-diverse regions harbor less genomic heritability for hermaphrodite self-fertility (Figure 2G), a known fitness-proxy in our environmental conditions [60]. Hence, the *rec-1* mutant tends to decrease recombination in genomic regions harboring fitness variation.

**Fig. 2.**
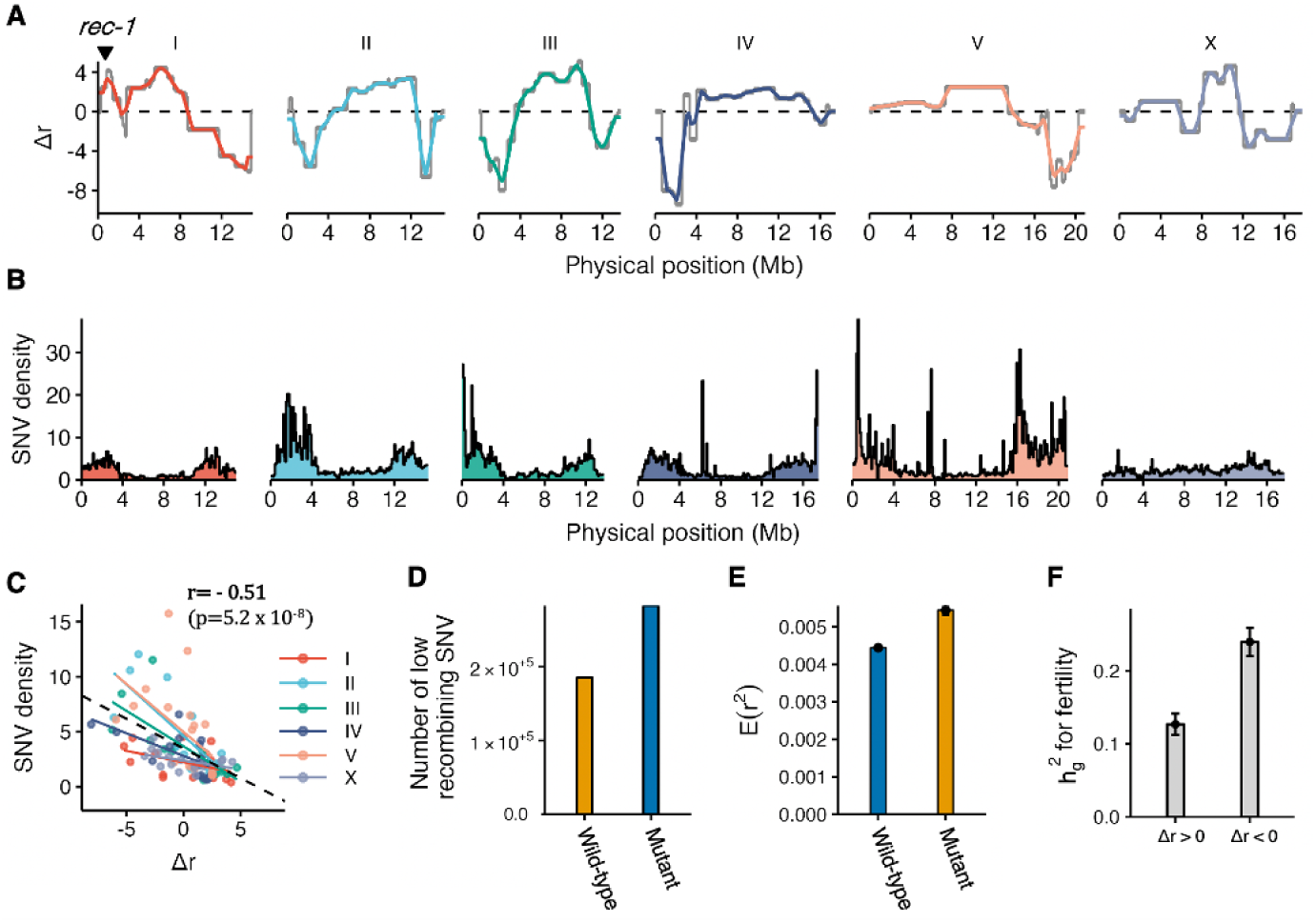
Genetic variance for fitness correlates with *rec-1* recombination landscapes. **A.** The grey line indicates the difference in recombination rates (cM/Mb) between the *rec-1* mutant and wild-type alleles (Δ*r*), estimated from the *rec-1* genetic linkage maps of [56] (see Methods). The colored lines indicate the Δ*r* calculated using the genetic distances corresponding to a 1Mb window around that position. In subsequent analyses, these colored lines define genomic positions where the *rec-1* mutant allele increases recombination relative to the wild-type allele (Δ*r >* 0) or reduces it (Δ*r <* 0). **B.** Density of SNV (number/kb) in 10^5^ bp non-overlapping windows. **C.** SNV density as a function of the difference of recombination rate between *rec-1* wild-type and mutant (Δ*r*) within 1Mb non-overlapping windows. Pearson correlation coefficient and p-value are shown. **D.** Proportion of SNV pairs having a probability of recombining lower than 1% (i.e., genetic distance *<* 1 cM) under the wild-type and mutant recombination landscapes among all possible pairs of 5,000 sampled SNV positions. **E.** For each SNV pair, the wild-type and mutant recombination probabilities are used to estimate the expected linkage disequilibrium, *E*(*r*^2^), at recombination - drift equilibrium (cf. [62]). Bars and errors are the mean and 95% CI for 1,000 random samples. **F.** Genomic heritability (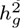; see Methods) for hermaphrodite self-fertility in the CeMEE panel, a collection of inbred lines sampling the diversity of the A6140 domesticated population [60, 61], for genomic regions where the *rec-1* mutant allele increases or decreases recombination relative to the wild-type allele.

### The mutant recombination landscape limits adaptation

By reducing crossovers in genomic regions that are genetically diverse and with more genetic variance for fitness, the *rec-1* mutant recombination landscape is expected to impede adaptation. To study adaptive dynamics under alternative *rec-1* recombination landscapes, we performed evolution experiments with the sister populations, fixed for either the *rec-1* wild-type or mutant allele (Figure S1). 24 replicate populations were cultured in the domestication or the novel salt environments at *N* = 10^4^ (Table S4). During 40 generations, outcrossing rates were high, independently of environmental conditions (Figure S5), except in two populations where self-fertilization became predominant and which were excluded from subsequent analysis as self-fertilization changes effective recombination rates (Figure S5). A third population was excluded from genomic analysis after facing the invasion of an obligately-selfing mutant (Figure S6; Figure S7)

Genome re-sequencing of samples taken at several time points during the 40 generations of experimental evolution indicates highly polygenic differentiation between populations at common SNV between the domestication and salt environments (Figure 3A; Table S4; Methods), but also within each environment (Figure S8). Consistent SNV differentiation among replicate populations was independent of *rec-1* allele identity. However, we find that the correlation between pairwise SNV differentiation, indicative of linked alleles, is higher among the *rec-1* mutant populations at the genomic regions where the mutant allele reduces recombination rates (Figure 3B; Methods). In contrast, the correlation between SNV differentiation is lower at the genomic regions where it increases recombination rates compared to the *rec-1* wild-type populations. As expected from the results of Figure 2E, a similar pattern is also found at the genome-wide level because most genetic diversity is found in regions where the mutant decreases recombination rates, increasing thus the genome-wide linkage. Therefore, these findings align with reduced recombination rates that potentially increase interference between selected alleles at many small-effect loci across the genome [36].

**Fig. 3.**
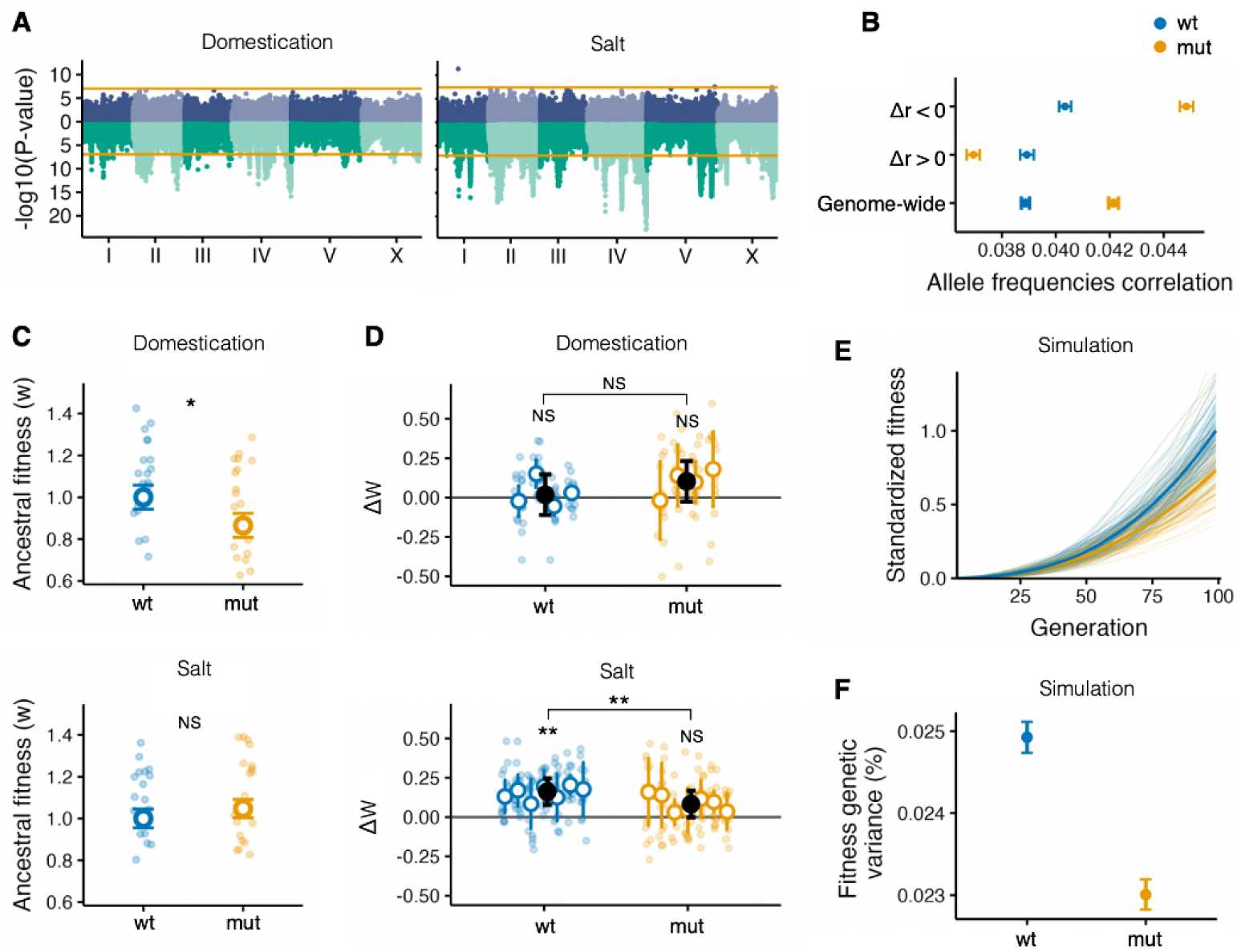
The *rec-1* mutant recombination landscape impairs adaptation. **A.** Consistent SNV differentiation among replicate populations during experimental evolution across the genome (green; GLM, Welch t-test on SNV allele frequency change with generation) and their dependency on *rec-1* allele identity (purple; GLM, Welch t-test on generation:rec1 interaction). The orange lines indicate the *α* = 0.05 threshold obtained by 1,000 random permutations of generation within a replicate population. The sampled populations and time points are in Table S4. **B.** Correlated SNV differentiation, calculated as the average pairwise Pearson correlation of SNV frequency change (1,000 SNVs sliding windows) among wild-type (blue) or mutant (orange) populations (both environments). This metric is calculated for SNV located within genomic regions where recombination is enhanced (Δ*r >* 0) or diminished (Δ*r <* 0) by the *rec-1* mutant, or genome-wide (1Mb window; Figure 2b). Error bars show the 95% CI obtained through 1,000 bootstraps of SNV. **C.** Fitness (w) of the *rec-1* wild-type and mutant ancestral sister populations (HR0 and MR0), obtained through competitions with a common tester strain and standardized on the wild-type sister population fitness (*w_wild−type_*=1) (see Methods). **D.** Adaptation (Δ*w*) of replicate populations after 40 generations of experimental evolution, calculated as the relative fitness difference (in %) from their respective ancestor population (HR0 or MR0). In **C,D.**, colored-filled dots are the technical replicate observations, and colored circles are the replicate population least-square estimates. Black dots and error bars indicate the least-square estimates of the *rec-1* wild-type and mutant mean and 95 % CI. **E,F.** For each *rec-1* allele, the evolution of 1,000 populations was simulated with the same setup as in Figure 3, but explicitly modeling *rec-1* mutant or wild-type alleles altering the recombination landscape. **E.** The average population fitness, standardized to yield 0 and 1 at generation 1 and 100, has a slower increase in the mutant populations (dark orange) than the wild-type population (dark blue). Light lines show individual replicate simulated populations. **F.** Restricted adaptation results from increased negative linkage disequilibrium during the simulations as mutant populations show less genetic variance for fitness (see Methods).

The effect of the recombination landscape on adaptation was assessed by measuring the fitness of each experimental population when competing with a morphologically-marked tester (*w*; see Methods; Table S6). In the domestication environment, the sister ancestral population fixed for the *rec-1* mutant allele shows reduced fitness when compared with the *rec-1* wild-type sister ancestor population (Figure 3C; Linear Model, Welch t-test on *rec-1* allele effect p-value=1.0 *×* 10*^−^*^3^). Genomic regions where the *rec-1* mutant increases recombination probably underlie this recombination load [27, 28, 33], and should contain beneficial epistatic genotype combinations selected during the 140 generations of domestication, as previously shown [58, 60]. In contrast, there was no opportunity for selection of epistatic genetic combinations specific to the salt environment in which the sister populations were never exposed prior to the experiment, and thus, they initially show similar fitnesses (Figure 3D; *rec-1* allele effect p-value=0.11).

Alternative *rec-1* recombination landscapes are inconsequential to further 40 generations of adaptation in the domestication environment (Figure 3D; Δ*w_wild−type_*=1.78 %; Δ*w_mutant_*=10.26 %: Linear Mixed Model, Welch t-test on *rec-1* allele effect *p − value* = 0.12; see Methods). As expected, however, in the novel salt environment, the *rec-1* mutant populations show less adaptation after 40 generations than *rec-1* wild-type populations when compared to their respective ancestor populations before experimental evolution (Δ*w_wild−type_*=16.21 %; Δ*w_mutant_*=8.28 %; Welch t-test on *rec-1* allele effect *p − value* = 2.5 *×* 10*^−^*^3^).

Individual-based computer simulations mimicking the demographic and genomic conditions of experimental evolution recover the impaired adaptive dynamics observed in the mutant populations (Figure 3E; see Methods). In the simulations, the lack of adaptation of *rec-1* mutant populations during experimental evolution is caused by a reduced disruption of negative linkage disequilibrium, diminishing the fitness variance when compared to *rec-1* wild-type populations (Figure 3F). Evolution under the *rec-1* mutant recombination landscape thus increases selective interference by reducing recombination in the genomic regions containing the relevant genetic diversity for polygenic adaptation.

### Hill-Robertson effects underlie local indirect selection on *rec-1*

To test for indirect selection on *rec-1*, we combined the two genetically diverse ancestor sister populations at equal ratios and performed experimental evolution (Figure S1). Because genetic diversity is present in this mixed population, recombination effectively generates novel genotype combinations during experimental evolution that can become associated with one of the *rec-1* alleles. Indirect selection coefficients (*s_i_*) were estimated from *rec-1* allele frequencies during 14-16 generations in either the domestication environment or the novel salt environment at *N* = 10^4^ across 20 replicate populations (Table S8). We find indirect selection for the *rec-1* mutant allele in both the domestication and novel environments (average between environments *s_i_*=6.75 *×* 10*^−^*^3^; linear model, Welch t-test p-value=4.1 *×* 10*^−^*^4^; Figure 4A), though indistinguishably between them (p-value=0.54).

**Fig. 4.**
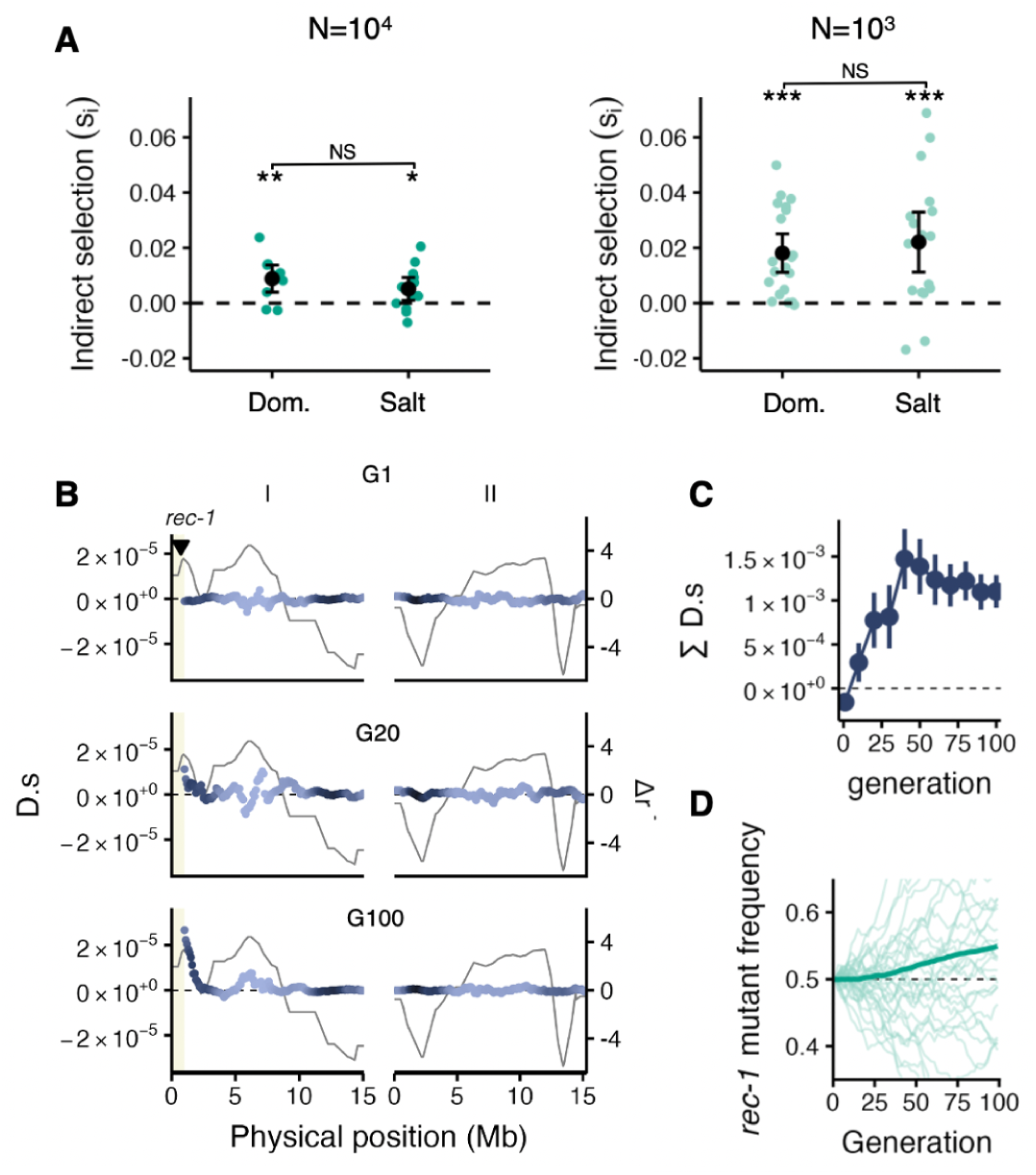
Indirect selection on the *rec-1* mutant is due to local associations with beneficial genotypes. **A.** Indirect selection coefficient (*s_i_*) of the *rec-1* mutant allele inferred from its change in frequency during 14-16 generations of experimental evolution in *rec-1* polymorphic populations with varying population size, in the domestication and salt environments. Each green dot is an independent replicate population (total n= 60). Black dots and error bars are the mean and standard error of the selection coefficients obtained by GLM. **B,C,D.** 1,000 simulation runs of chromosomes I and II from 200 mimics of ancestral populations (5 runs per ancestral population) obtained by sampling 1,000 SNV of the domesticated A6140 population under a multiplicative fitness model (see Methods). To simulate the experimental loss of diversity around *rec-1* due to the introgression into the domesticated A6140 population, no SNV was sampled in the first 1Mb of chromosome I. **B.** The associations of the *rec-1* mutant allele with beneficial alleles generated during simulations explain indirect selection at the *rec-1* locus. These associations are highlighted by multiplying the linkage disequilibrium between each selected allele and the *rec-1* mutant allele (D; [63]) by its selection coefficient (as the strength of indirect selection at the *rec-1* locus depends on this product, [21]). The quantity *D · s* (purple line) is positive when the *rec-1* mutant allele is associated with the favored alleles. Purple lines indicate *D · s* when averaged over 1Mb sliding windows, with lighter shades showing lower SNV density (Figure 2c). The grey lines indicate the difference in recombination rates due to *rec-1* loss of function (Δ*r* from Figure 2b). The triangle indicates the *rec-1* locus. At the first generation of the simulations (G1), the *rec-1* mutant allele is not associated with selected alleles. During evolution (G20, G100), it generates associations with beneficial alleles (*D · s >* 0) in its vicinity by breaking local negative LD. **C.** Average sum of all the associations (Σ*D · s*) across the two chromosomes is largely determined by the genomic region around *rec-1*. Detailed analysis of how the modifier becomes associated with surrounding genotypes along chromosome I is shown in Figure S11 and Figure S12. **D.** *rec-1* mutant allele frequency during the simulations (dark green is the average among replicates).

Simulations of experimental evolution, and assuming no direct selection at the *rec-1* locus (see Methods) show that the indirect selection that is observed can result from the associations *rec-1* alleles established with recombinant genotypes located in their vicinity (Figure 4B,C,D; Figure S11, Figure S12). While the *rec-1* mutant alters recombination on all chromosomes, a modifier can only be indirectly selected with the genetic combination it creates nearby, as it can not maintain associations with more distant alleles [24, 44]. By locally increasing recombination rates on the left side of chromosome I (cf. Figure 2A), the *rec-1* mutant allele is under indirect selection because it breaks negative linkage disequilibrium at the fitness loci surrounding it, hitchhiking then with the beneficial local genotype it creates. The size of the genomic region explaining indirect selection during simulations is not usually larger than 3 Mb on the right side of the *rec-1* locus. However, transient beneficial associations can be generated up to 7 Mb (Figure S11, Figure S12), despite the low genetic diversity in the chromosomal central regions.

Population genomic data from the fixed recombination landscape experiments (cf. Figure 3) show that SNV responses within the region surrounding *rec-1* do not differ between environments (Figure S8), thus explaining why indirect selection is similar between environments. Nevertheless, estimates from the fixed recombination landscape experiments also indicate that the haplotype mass-introgressed together with the *rec-1* mutant has a small positive selection coefficient (*s_h_*) of similar magnitude than *s_i_* (domestication: *s_h_* = 3.6 *×* 10*^−^*^3^; salt: *s_h_*= 1.1 *×* 10*^−^*^2^; Figure S9; Methods). This raises the possibility that the *rec-1* mutant is favored because it is initially linked with beneficial alleles in the ancestral population rather than indirect selection resulting from the association with the adaptive genetic combinations it creates during experimental evolution.

Indirect selection is expected to be correlated with the extent of genetic drift because negative linkage disequilibrium is generated through the stochastic effect of interference between selected loci (Hill-Robertson effect) [30–32]. We performed experimental evolution of 40 mixed sister replicate populations with reduced size of *N* = 10^3^ (Table S8) in both the domestication and salt environments. We find that at this smaller population size, indirect selection on the *rec-1* mutant allele is greatly enhanced when populations evolve at *N* = 10^3^ (Figure 4a; *s_i_*=2 *×* 10*^−^*^2^, Linear Mixed Model, Welch t-test for population size effect p-value=3.19 *×* 10*^−^*^3^). Simulations indicate that many of the associations that explain selection on the *rec-1* mutant are expected to be transient (Figure S12), and genome re-sequencing of wild-type and mutant isogenic lines obtained from evolved salt small populations (Methods; Table S2) does not highlight persistent associations between the *rec-1* alleles and recombinant genotypes generated during experimental evolution (Figure S10). Overall, these findings indicate that indirect selection on recombination is due to its effect in breaking negative linkage disequilibrium created during experimental evolution by the Hill-Robertson effect.

### Conclusions

Our experimental system has allowed us to validate several theoretical expectations on the evolution of meiotic recombination rates in already-recombining populations [20–23, 25]. Increasing recombination in genomic regions harboring more genetic variation for fitness enhances adaptation, but this does not necessarily translate into selection for modifier alleles having such an effect. It has been recognized for long that indirect selection on recombination modifiers may not favor recombination rates that increase adaptation and population mean fitness, particularly in the presence of epistatic interactions between fitness loci [21, 64]. Our results pinpoint another possible explanation based on contrasted local and global effects of recombination landscape modifiers. Indeed, indirect selection primarily depends on genetic associations between the modifier and nearby selected loci that are maintained over a higher number of generations than associations with more distant loci [21, 24, 44]. In particular, a modifier allele increasing the efficiency of selection in its genomic neighborhood (by locally increasing recombination) may be favored even though its overall genomic effect is to reduce recombination between fitness loci. We can conclude, therefore, that under heterogeneous recombination landscapes, indirect selection for recombination is decoupled from polygenic adaptation.

Our findings help reframe several questions about the evolution of recombination, maintenance of standing genetic variation, and adaptation in natural populations. While modifiers increasing the density of crossovers uniformly along chromosomes should generally favor adaptation, we predict that modifiers affecting the positions of crossovers should have more contrasted effects, the consequences of the evolution of recombination on adaptation depending strongly on the relative contribution of their genomic neighborhood to the variance in fitness. The size of the relevant genomic region is expected to increase as the fitness effect of selected loci increases and as recombination rates decrease [21, 24]. In particular, the fate of recombination modifiers should depend more on their overall genomic effect when effective recombination rates are reduced by inbreeding or population subdivision [48, 65–68]. Ultimately, the presence of multiple recombination modifiers segregating within populations, as observed in natural populations including in humans and species of agricultural and conservation significance [69], opens the possibility for conflicts among those modifiers, which may lead to rapid evolution of recombination rates and interact with local genomic features such as chromatin structure, gene or transposon density [6–8, 70]. Addressing these and other outstanding questions will require developing new theoretical and empirical models of polygenic adaptation under variably heterogeneous recombination landscapes.

## Methods

### C. elegans culture

All *C. elegans* culture protocols followed ref. [57]. Experimental population samples were revived from −80°C cryopreserved stocks with at least 10^3^ individuals and expanded for two generations in a common environment before use. In the standard domestication environment, populations were cultured in 9 cm Petri dishes filled with NGM-lite agar (US Biological) containing 25 mM NaCl and a 100 µL lawn of *E. coli* (HT115) as a food source for growth until reproduction at 20°C and 80% RH. In the novel high-salt environment, 230 mM of NaCl was added to the NGM-lite agar. Populations were passaged under a 4-day non-overlapping life-history stage cycle. At each generation, synchronized L1 starved larvae were seeded onto Petri plates at a density of 10^3^ individuals per plate. Following 72 hours of growth and reproduction, adults and embryos were collected using M9 (22mM KH2PO4, 42mM Na2HPO4, 85mM NaCl, and 1mM MgSO4) and exposed to 20mM KOH:0.6% NaClO for 5 minutes. This “bleach/hatch-off” treatment results in adult and larval death while allowing embryo survival. After three washes with M9, the embryos were incubated in M9 at 20°C with aeration for 24 hours. Live L1 densities were estimated by scoring their number in 15-50 *µl* of M9 under a Nikon SM1500 dissecting scope. Pooled samples of individuals for DNA analysis and/or cryogenesis were collected 24 hours after the bleach/hatch-off treatment. They all contained *>* 10^3^ live L1s with larval and adult debris.

### Experimental evolution

Three sets of evolution experiments were performed. Figure S1 shows a schematic of the experimental designs, Table S1 a general description of the base populations, and finally Table S3, Table S8, and Table S4 detailed lists of replicates and sampling for population genomics and/or adaptation assays of each set of evolution experiments.

In the first set of evolution experiments (Experiment I; Table S3), we tested for direct selection (*s_d_*) on *rec-1* in populations that differed only in *rec-1*allele identity; see [56] for the derivation of *rec-1* wild-type and loss-of-function mutant. For each population, the EEV1401 strain (*rec-1* wild-type) and the EEV1403 strain (*rec-1* mutant in EEV1401 genetic background), or the EEV1402 strain (wild-type) and the EEV1404 strain (mutant, EEV1402 background) were combined into a single population. A total of 143 replicate populations were then maintained in one Petri dish each (*N* = 10^3^ L1s) for five generations while varying isogenic background, initial sex ratio, initial *rec-1* allele frequency, outcrossing rate, or salt environment. The initial sex ratio was manipulated by hand-picking 50 immature L4 hermaphrodites and 50 males for the high male treatment or 50 immature hermaphrodites for the low male frequency treatment two generations before the start of the experiments. *rec-1* allele frequencies were manipulated by mixing EEV1401 (EEV1402) with EEV1403 (EEV1404) at 1:1 or 4:1 ratio at the setup of the experiments (G0 generation). Samples were collected at generations 1, 3, and 5 for *rec-1* genotyping.

The second set of evolution experiments tested for adaptation under alternative *rec-1* recombination landscapes (Experiment II), while the third set for indirect selection (*s_i_*) at *rec-1* (Experiment III). Both sets of experiments relied on the derivation of two ancestral sister populations recovering most of the standing genetic variation of the A6140 domesticated population [59, 60] but differing in *rec-1* allele homozygosity (Table S1). These sister *rec-1* wild-type and *rec-1* mutant populations are denoted as HR0 (for Heterogeneous Recombination, generation 0) and LR0 (Leveled Recombination, generation 0), respectively.

HR0 and LR0 were obtained by crossing males from the A6140 population with the *rec-1* mutant EEV1403 strain. A total of 340 successful crosses were conducted. From each cross, one F1 immature hermaphrodite was isolated, and 12 F2 hermaphrodites were allowed to self individually. Pooled F3 progeny samples were collected for *rec-1* genotyping (see below). From each F2 family, one F3 *rec-1* homozygous wild-type sample and one F3 *rec-1* homozygous mutant sample were expanded and cryopreserved. In total, we derived 340 wild-type and 340 mutant samples. To obtain the HR0 and LR0 populations, we first combined L1s in equal proportions and, to increase the frequency of males, adults were harvested 96 hours after and filtered using a 0.11 *µ*m nylon filter (Merck NY1100010) to separate males from hermaphrodites, remixed at a higher male proportion, and allowed to mate for 24h. The next generation followed a similar procedure. The population sizes were above *N* = 10^4^ during these two generations. HR0 and LR0 are the ancestor populations for the experiments on adaptation under alternative *rec-1* recombination landscapes. We cultured four and eight replicate populations, from each HR0 and LH0, in the domestication and novel salt environments, respectively, for 40 generations and at *N* = 10^4^ L1s (10 Petri dishes). Pooled samples of individuals from 16 replicate populations at generations 0, 6, 17, 24, 30, 33, and 40 were collected for genome re-sequencing (Experiment II; Table S4).

To test for indirect selection, we first established the MR population (Mixed Recombination) by combining HR0 and LH0 at an equal L1 ratio. Next, and to break the linkage between the *rec-1* mutant allele and the EEV1403 introgressed allele, the MR population was maintained at *N* = 10^5^ for five generations. The ancestral MR0 population (MR, generation 0) was obtained after these five generations. A total of 19 replicate populations were evolved at *N* = 10^4^ and 41 replicate populations at *N* = 10^3^ (SMR#, for Small size MR replicate number) during 14-16 generations in the domestication and salt environments (Experiment III; Table S8). Different populations were cultured at different calendar dates (blocks). From SMR24, SMR25, and SMR40 replicates at generation 14, we sampled immature hermaphrodites and inbred these lines by selfing for 10 generations. The resulting inbred lines were subsequently genotyped for *rec-1* and genome re-sequenced.

### DNA extraction

DNA was extracted from samples with adult debris following the bleach/hatch-off protocol. The collected worm material was incubated in 600 *µ*L of Cell Lysis Solution (Qiagen ref. 158906) with 5 *µ*L of proteinase K (20mg/ml) overnight at 56°C with 700 rpm shaking. This was followed by a 1-hour incubation at 37°C with 10 *µ*L of RNAse A (20mg/ml) (Roche). Subsequently, the lysates were cooled on ice and mixed with 200 *µ*L of Protein Precipitation Solution (Qiagen ref. 158910) for 5 minutes. After centrifuging at 15000 rcf at 4°C for 10 minutes, the supernatant and 600 *µ*L of isopropanol (VWR chemicals) were mixed by inversion and incubated for 1 hour at

-20°C. Precipitated DNA was pelleted with a 10-minute centrifugation at 15000 rcf and 4°C and pellets washed with 600 *µ*L of 70% ethanol, air dried for 15 minutes, and resuspended in 50 to 200 *µ*L of Tris-EDTA buffer (Fisher Bioreagents, BP2473-100). DNA quantification was done using a Qubit spectrophotometer (Invitrogen) with a dsDNA BR assay kit (Invitrogen Q32853). To achieve the desired concentration, samples were diluted in DNAse-free water (Invitrogen, 10977-035).

### *rec-1* pool-genotyping

We quantified *rec-1* mutant allele frequencies (*p*) in samples of pooled individuals from each population (Experiment I and III) or from families (HR0 and MR0 derivation) by developing a new protocol using quantitative PCR (qPCR; Figure S13, details in our Github). In brief, during the melting phase following the amplification step of qPCR, the temperature at which double-stranded DNA (dsDNA) transitions to single-stranded DNA (ssDNA) depends on the sequence and length of the DNA fragment. This property can identify different DNA sequences by analyzing the “melting” profile of the qPCR [71, 72]. For this, 15 ng of DNA was amplified in 15 *µ*L reactions containing 7.5 *µ*L of SYBR Green Master mix (Roche, 04887352001) and 0.03 *µ*M of primers [forward: GCAGGTTTTAGCAGAAAAACG; reverse: CGTCGGAACGTATCCTGGT;[55]; Eurofins Genomics] during 26 to 35 cycles of temperature steps alternating between 95°C and 60°C using a LightCycler 480 (Roche Diagnostics). Amplification yields *rec-1* wild-type and mutant alleles of 71 bp and 63 bp, respectively, as the *rec-1* mutation used here is an 8 nucleotide deletion [56]. Following amplification, melting was initiated, gradually raising the temperature from 60°C to 95°C at a rate of 0.02°C/s with 25 data acquisitions per °C. The melting data were extracted using LightCycler 480 1.5.1 software and analyzed using R [73]. After exponential background subtraction (following [71]), analysis of the normalized melting profile enables the achievement of accurate quantification of the relative frequency of *rec-1* mutant alleles (*p*) within a sample. Calibration samples were included for each qPCR plate run by mixing EEV1401 (wild-type) and EEV1403 (mutant) samples with known mutant allele proportions: 0%, 10%, 25%, 50%, 75%, 90%, and 100% (see Figure S13 for details).

### Selection coefficients

Selection coefficients were defined as 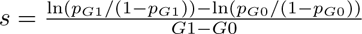, where *G* is the generation and *p* is the *rec-1* mutant allele frequency [74]. They were estimated for each replicate population in experiment I (*s_d_*; direct selection) or experiment III (*s_i_*; indirect selection) by fitting a Generalized Linear Model (GLM) to the logit-transformed *rec-1* mutant allele frequency (*p*) as a function of generation (in R syntax: *p ∼ generation*, the error family set to ‘binomial’). Selection coefficients at variants other than at *rec-1* are similarly defined in the population genomic analyses of experiment II (*s_h_*) and simulations (see below).

For *s_d_* estimates shown in Figure 1b, the “global” effects were estimated by fitting *p ∼ generation* across all replicates. The effects of the several treatments (environment, genotype, and sex ratio) were modeled separately (*p ∼ generation ∗ treatment*). A single GLM with all treatments and accounting for pseudo-replication (indicating block structure using the glmer function; [75]) failed to converge due to the uneven number of replicates within each Table S3. Hence, to account for pseudo-replication, the 95% credible interval estimates were obtained by 10^4^ bootstrapping with resampling replicate populations, with the p-value determined by the proportion of the bootstrapped distribution overlapping zero for each GLM.

Testing of *s_i_* ≠ 0 in the small population size experiment (*N* = 10^3^) used Welch t-tests on Linear Model intercepts (LM; *s_i_ ∼* 1). To test for treatment effects a Linear Mixed Model was fitted (LMM; *s_i_ ∼* (1*|block*) + *environment* + *N*), where block is a random factor reflecting when the experiments were done (Table S8), environment for domestication or salt fixed effects, and *N* for the population size fixed effects. REML estimates were tested for significance with Welch t-tests.

### Genome re-sequencing

Library preparation and sequencing were conducted at BGI Hong Kong Tech Solution (random library selection; BGISEQ-500; 150bp paired reads) at 10X and 60X coverage for the SMR inbred lines and the experimental populations. The FASTQ files were aligned to the WS245 *C. elegans* reference genome [76] using *BWA* [77], allowing for up to two mismatches. Paired reads where both reads were not correctly mapped or with a low mapping quality score were filtered out (samtools view; -q 20 -f 0×0002 -F 0×0004 -F 0×0008 options; [78]). A total of 371,130 single nucleotide variants (SNV) that were known to segregate in the A6140 ancestor of these populations [61] were called using *freebayes* [-@ option; [79]]. The 5% of variants with the smallest coverage depth were filtered out, leaving 367,917 SNV available for population genomic and heritability analysis. Sequence read archives will be available in NCBI (inbred lines: PRJNA1047336; populations: XXXXXX) and the genotype tables for the called SNV in GitHub.

### Population genomics

Using *rec-1*genetic linkage maps from ref. [56], the mutant minus the wild-type recombination rates (cM/Mb) between two genomic positions, referred to as Δ*r*, was calculated. Δ*r* and SNV density (SNV number per kb) were estimated within 1Mb non-overlapping windows (Figure 2C). Furthermore, for each SNV, Δ*r* was calculated in 1Mb sliding windows between the positions 500 kb before and after their genomic position (colored lines in Figure 2A) to define whether they were located in a region where the *rec-1* mutant increases (Δ*r >* 0) or decreases (Δ*r <* 0) the recombination rate relative to the wild-type. We estimated the expected average pairwise linkage disequilibrium between SNV at drift-recombination equilibrium as 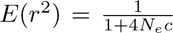, following ref. [62], where *c* is the *rec-1* wild-type or mutant genetic distance between SNV, from ref. [56], and *N_e_* = 10^3^, the effective population size, from ref. [58].

For each SNV across the genome, we estimated the deviation from 0 in the average allele frequency change for each SNV using a Generalized Linear Model (GLM) of SNV allele counts across generations (*SNV.count ∼ generation*; error family set to ‘quasibinomial’; glm function in R; [73]). The interaction effects between generation and the *rec-1* allele identity were estimated using a similar GLM (*SNV.count ∼ generation ∗ rec*1; Welch t-tests for the significance of *generation*: *rec*1 interactions). A separate set of GLM models was used to identify the interaction of generation with the environment (*SNV.count ∼ generation ∗ environment*; Welch t-test on *generation*: *environment* interaction). Significance thresholds were estimated by generating 10^3^ permutations of generation identity within each population. The minimum p-value within each permuted dataset was recorded. The significance threshold corresponds to the 5th percentile of the distribution of these minimal p-values.

The pairwise Pearson correlation was calculated for each possible pair of SNV within 1,000 SNV non-overlapping windows for the wild-type or mutant samples. By pooling the samples from the domestication and salt environments, the replication and the differentiation of allele frequencies were maximized to increase the power to detect the effect of recombination on this correlation. Pairwise correlations were averaged for the SNV found within regions where the *rec-1* mutant increases recombination (Δ*r >* 0; 1Mb sliding window; see Figure 2A), or where it reduces recombination (Δ*r <* 0). 95% credible intervals for the average pairwise correlations were obtained by resampling 10^3^ times all SNV used to calculate the correlation.

Inbred line “founder haplotype blocks” were assessed on chromosome I by phasing founder genotypes following ref.[60]. Briefly, the homozygous genotype of a given inbred line is matched with the homozygous genotype of the founders at multiple overlapping windows of varying sizes. The different windows are then extended until the inbred lines’ genotype does not match a single founder genotype (or a single set of founders when they share a common genotype), indicating the window reaches a haplotype block boundary. Redundant haplotype information was filtered out. Details of the methods and R scripts can be found in our GitHub repository.

### Genomic heritability

Genomic SNV heritability 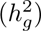 for hermaphrodite self-fertility was estimated using 240 genotyped inbred lines that had been phenotyped in the domestication environment as part of the CeMEE panel, a set of recombination inbred lines derived from the domesticated A6140 population and related populations [60]. Heritability is obtained as 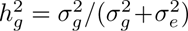, where 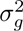 is the additive genetic variance and 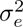 the environmental variance [74]. Genomic heritability was estimated from linear mixed-effects models assuming that SNV random effects followed normal distributions centered on zero and with variance 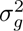 times a genetic similarity matrix *A* accounting for relatedness of the inbred lines [80]. The residual variance of the models is 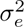. To estimate *A*, we followed ref. [81] and assumed that each SNV explains the same amount of genetic variance, i.e., that an inverse relationship exists between SNV effect size and allele frequency. Misestimation of 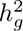 can occur because SNVs inevitably tag the (unknown) causal alleles to varying extents due to differences in linkage disequilibrium (LD). SNVs in high LD (*r*^2^ *>* 0.9) were therefore pruned, resulting in a 40,779 SNV dataset (see Table S5). We fitted one model with all genome-wide SNV and another with SNV from genomic regions with Δ*r >* 0 and Δ*r <* 0 as independent random effect factors. Models were fitted with the *mmer* function from the *sommer* package in R [82]. Scripts for this analysis can be found in our GitHub repository.

### Adaptation assays

Fitness was quantified in one-generation competitions of the experimental populations against the EEV1402 strain carrying a visible green fluorescent protein (GFP) construct (Table S1). A schematic of the assays is presented in Figure S14 and follows the design of ref. [59]. Fitness of an experimental population was defined as the proportion of wild-type alleles transmitted after one generation of competition relative to the GFP tester: *w* = 1 + ln(*p_wt.g_*_1_*/p_GF P.g_*_1_)-ln(*p_wt.g_*_0_*/p_GF P.g_*_0_); *p_wt_* and *p_GF P_* being the wild-type non-GFP and GFP estimated allele frequencies, respectively, before (*g*0), and after the competition (*g*1). For the populations after 40 generations of experimental evolution, the percentage change in adaptation (Δ*w*) is given by their fitness divided by the average fitness estimates of their respective ancestor population when assayed in the same calendar date and the same environment. Because GFP is dominant, heterozygosity must be inferred from the proportion of the GFP alleles after one generation of competition. GFP heterozygosity was estimated using the mating Table S7 under the assumptions of similar self-fertilization rates and no assortative mating between tester strain and experimental population.

To conduct the assays,10^3^ individuals were first thawed from −80°C stocks and synchronized at the L1 stage through the bleach/hatch-off protocol. In domestication or salt environments, an additional generation was cultured with 7 *×* 10^3^ individuals. In parallel, the EEV1402 tester strain was thawed and maintained in the domestication environment for two generations. In the third generation post-thawing of samples, the assays were set up by mixing synchronized L1s from the experimental populations and the tester strain at a 1:1 ratio at a density of *N* = 10^3^ in the domestication or the salt environments. The proportion of GFP vs non-GFP individuals was scored under a dissection microscope in the initial L1 mix (*g*0) and after one generation at the L1 stage (*g*1), by counting individuals on a glass slide (median = 403 individuals per competition replicate). Assays were conducted in four experimental blocks, each corresponding to independent thawing and culture calendar dates. Within each block, competitions for each evolved population at generation 40 of experimental evolution were replicated three to five times, resulting in 11 to 16 technical replicates per population (mean = 15.4). Competitions for ancestral populations (HR0 and LR0) were replicated three to six times within each block, resulting in 21 replicates each (Supplementary Table S6). For each environment independently, adaptive differences between the two *rec-1* ancestral populations (HR0 and LR0) were assessed by fitting a Linear Mixed Model (LMM; *lme4* R package; [75]): *w ∼ ancestral* + (1*|block*); with ancestral as a fixed effects factor and block as a random effects factor. The significance of ancestral adaptive differences was assessed with Welch’s t-test. The average level of adaptation after experimental evolution of the wild-type or evolved population was assessed by testing if Δ*w* is different from zero using LMM: Δ*w ∼* 1 + (1*|block*) + (1*|population*); where population was the replicate population. Welch’s t-test was used to test for a significant intercept. Finally, the adaptive differences between wild-type and mutant evolved populations (*rec*1) were modeled in as: %Δ*w ∼ rec*1 +(1*|block*) +(1*|population*); with Welch’s t-test employed to test for significance of *rec*1 fixed effects. Models were fit with REML, and confidence intervals were computed with the *emmeans* R package [83].

### Males and hermaphrodites

The consequences of NaCl for hermaphrodite self-fertility and male frequencies in the MR population were determined in the domestication environment (25 mM NaCl in the growth media) and at high concentrations (200 mM, 240 mM, 275 mM). Hermaphrodite fertility was estimated, following two generations of exposure to the focal NaCl environment, by counting the number of *in utero* eggs in adults, fixed following [84], under a Nikon AZ100 equipped with a camera. Male frequencies were estimated by scoring sexes after four generations of exposure to the focal NaCl environments. As hermaphrodites cannot mate with each other, and sex-chromosome segregation is Mendelian, outcrossing rates can be estimated as twice the male proportion in the previous generation [85].

During Experiment II, male frequencies were measured 72 hours post-L1 using the *Multi-Worm Tracker* software [MWT 1.3.0; [86]], and as described in ref. [87] (Figure S5). Briefly, six-minute movies were recorded for each sample, and the extracted MWT trait values (movement, size, etc.) – for up to 10^3^ features (individuals) – used for the training of an extreme gradient-boosting model [*xgboost* R package; [88]] and differentiate hermaphrodites and males (details in our GitHub). Male frequencies obtained through this method are similar to those obtained by scoring sexes under a dissection scope at few common time-points (Pearson correlation: domestication: *r* = 0.66, *p − value* = 6.797 *×* 10*^−^*^6^; salt: *r* = 0.74, *p − value* = 4.13 *×* 10*^−^*^15^). As expected, outcrossing were lower in salt (Welch’s t-test: *p*-value = 9.8 *×* 10*^−^*^7^), but remained elevated in both environments (domestication:0.77, salt:0.66). No difference was found between the *rec-1* wild-type and mutant populations. The SLR3 (mutant) and SHR4 (wild-type) populations exhibited lower male frequencies compared to the average among all populations (populations; LM: *outcrossing.rate ∼ population* + *generation*; Welch t-test on population effect) and were excluded from subsequent analysis (Figure S5; Table S4). One of these two populations was genome re-sequenced and found to be an outlier (Figure S6). During the first block of Experiment III, outcrossing rates obtained by visually scored male frequencies in 72h-post-L1 adults, were, lower in remained elevated in both environments (domestication:0.94, salt:0.85). During the second block of Experiment III, visual inspection at every generation indicated that male frequencies remained high in every population.

### Simulations

Individual-based simulations of experimental evolution were implemented using *SLiM* 4.0.1 [89]. The simulation scripts and results are available on our GitHub repository. We modeled androdiecious diploids with hermaphrodites capable of self-fertilization and of outcrossing only with males. At each life cycle, offspring genotypes resulted from selecting two parental gametes. The origin of the first gamete was randomly sampled among hermaphrodites weighted by their fitness. For offspring resulting from self-fertilization, the second gamete originates from the same parental hermaphrodite. In the case of offspring resulting from mating between hermaphrodites and males, the origin of the second gamete was sampled among males weighted by their fitness. The proportion of outcrossed offspring was fixed at 80% and population sizes at *N* = 10^3^ in all simulations.

Gametes were created by recombining the two parental haploid genomes, with each chromosome pair having a 50% probability of undergoing one recombination event, corresponding to a genetic linkage map length of 50 cM per chromosome. The probability of recombination at a given position was directly proportional to the known recombination rates of wild-type or mutant *rec-1* from ref. [56]. In simulations where both *rec-1* alleles segregate, a recessive recombination modifier locus was explicitly modeled at the *rec-1* genomic position. A6140 CeMEE panel haplotypes [60] were simulated for a subset of 1,000 randomly sampled SNV distributed on chromosomes I and II. At setup, the initial population mimics the A6140 domesticated population (ancestor to all the populations employed during experimental evolution). Simulated selected alleles are thus expected to follow the experimental SNV density (Figure 2B), allele frequency spectrum (mostly extreme frequencies), and linkage disequilibrium (high under 0.5 cM of the wild-type maps). The selection coefficients were randomly assigned to one of the alleles by drawing from a normal distribution centered at zero. A large range of standard deviations in this normal distribution was implemented to interpolate the one yielding a genetic variance in fitness of the initial population of 0.03. Before simulations, negative LD, expected to be already present in these populations because of past selection, was generated through a burn-in of 100 generations under the wild-type recombination maps and negative epistasis (intermediate genotypes have higher fitness, extreme genotypes have lower fitness).

In simulations where both *rec-1* alleles segregate, we further sampled SNV based on the observed HR0 and MR0 SNV diversity to reflect the introgression of the EEV1403 allele. Individual fitness (*W*) in these simulated ancestral population mimics was defined following a multiplicative fitness model with codominance: 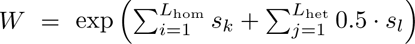; where *s* is a selection coefficient at the *k^th^* or *l^th^* of the *L*_hom_ homozygous or the *L*_het_ heterozygous loci.

Two hundred populations were generated from the A6140 ancestral mimic, from which five replicate simulations were run for 100 generations under fixed *rec-1* wild-type or mutant recombination rate landscapes, or where both *rec-1* alleles started at 1:1 proportions. The mean and variance in fitness of the simulated populations, as well as *rec-1* allele frequency, when relevant, were recorded. We further estimated the association of the *rec-1* mutant allele with beneficial alleles by calculating linkage disequilibrium (*D*; [90]) between them multiplied by their selection coefficient (*s*). *D · s* is positive (negative) when the *rec-1* mutant allele is associated with beneficial (deleterious) alleles.

## Data accessibility

Genome re-sequencing data will be deposited at NBCI (PRJNA1047336, XXXXX). All other data and code for analysis are available at our GitHub repository (see Table S2) and will be deposited in a public archive upon publication.

## Acknowledgments

We thank H. Gendrot, L. Gervasoni, J. Goņcalves, F. Mallard, V. Pereira, L. Sénéchal and R. Weissman for help with worm handling and data analysis. We thank D. Abu-Awad, M. Desai, C. Dillmann, C. Haag, T. Lenormand, F. Mallard, M. Rockman, R. Stetsenko and J. Yanowitz for discussion or comments on the manuscript. We have no competing interests to declare.

## Author Contributions

T.P.: Conceptualization, Data curation, Software, Formal analysis, Validation, Investigation, Funding acquisition, Writing; L.N.: Data curation, Formal analysis, Funding acquisition, Reviewing; D.R.: Conceptualization, Funding acquisition, Supervision, Reviewing; H.T.: Conceptualization, Resources, Supervision, Funding acquisition, Investigation, Writing, Project administration.

---

**Fig. S1.**
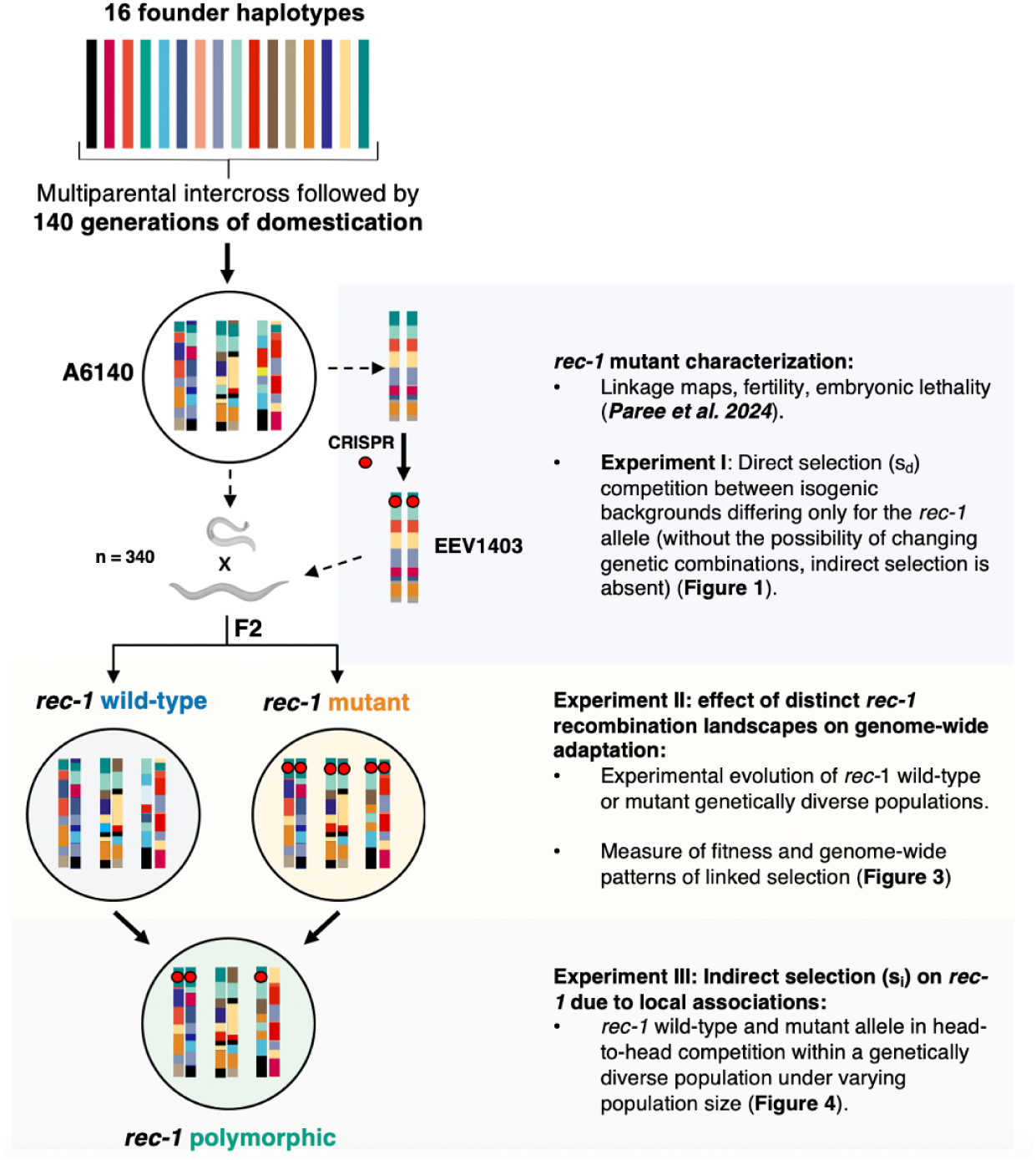
Experimental design. **A.** Populations and inbred lines were derived within the same framework to test the adaptive consequence and selection on the *rec-1* mutant. The base material for this study is the genetically diverse outcrossing A6140 population, which originates from the cross of 16 founders and has been adapted, during 140 generations, to a standard lab environment, referred to as the domestication environment [57, 60]. The EEV1403 line was generated by editing a loss-of-function mutation (CRISPR-cas9) in the *rec-1* gene in the EEV1401 inbred line [56], previously derived from the A6140 population [91]. The EEV1401/EEV1403 pair, among others (not shown), was used to construct *rec-1* linkage maps and study the effect of the mutant on fitness-related traits [56], as well as measure its direct selection coefficient (*s_d_*; Experiment I; Figure 1). Then, the EEV1403 line was crossed back to the A6140 population 340 times to generate two sister populations, wild-type or mutant, derived from same-family F2. The sister populations were then mixed in equal proportion to generate a *rec-1* polymorphic population, which was further maintained for 5 generations to break the linkage between the *rec-1* mutant and the EEV1403 introgression haplotype. The wild-type and mutant sister populations were used to address the adaptive consequence of alternative *rec-1* landscape (Experiment II; Figure 3). The *rec-1* polymorphic population was used to measure indirect selection (*s_i_*) on the *rec-1* mutant, occurring when there is genetic diversity and that recombination modifies the haplotype pool (Experiment III; Figure 4).

**Fig. S2.**
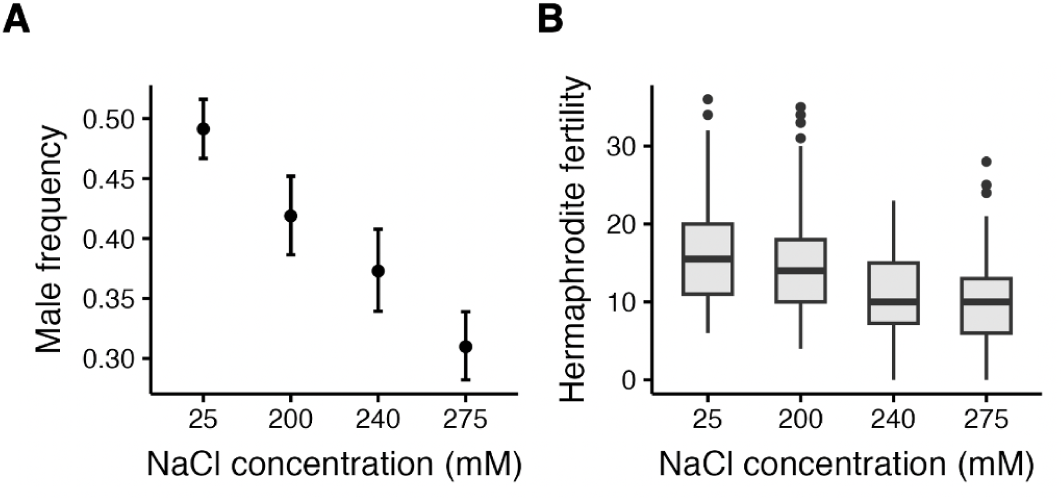
Exposure to salt concentrations in growth media impairs fitness proximal traits. The effect on male frequency and hermaphrodite fertility in MR population in the domestication NaCl (25 mM) and novel high NaCl concentration (200, 240, or 275 mM) environments. **A.** Male frequency after four generations of exposure to NaCl from an initially high male frequency (*∼*45%). As *C. elegans* hermaphrodites can only mate with males (and not each other), and because chromosomal sex determination has Mendelian segregation [85], the male frequency can be taken as a proxy of male mating ability during the four generations of the assay [59]. Points and error bars represent the least square estimates and 95% CI **B.** Hermaphrodite self-fertility is approximated by counting the *in utero* egg number in adults following two generations of exposure to NaCl (Methods). Each point represents a single hermaphrodite, with box plots showing the median and interquartile ranges for 451 hermaphrodites.

**Fig. S3.**
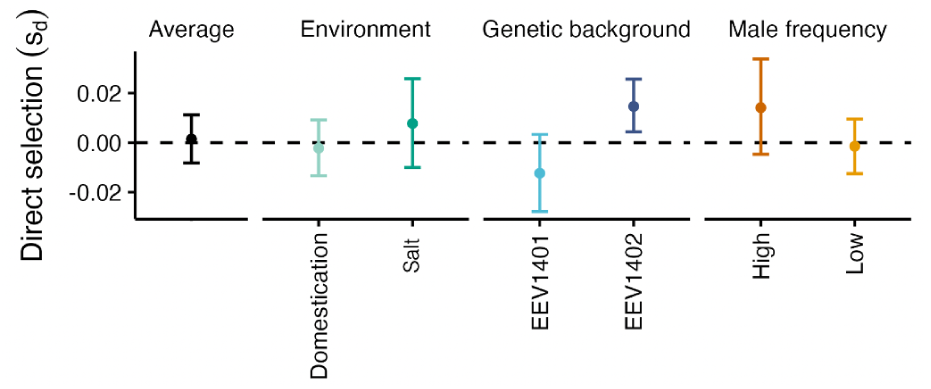
Effect of different treatments on direct selection. The black dot shows the average direct selection coefficient of the *rec-1* mutant (*s_d_*) inferred through GLM on all the replicate populations and already presented in Figure 1. The colored dots show the *s_d_* inferred for the variables. Error bars are the 95% credible intervals (CI) estimated by 10^4^ bootstraps with resampling of replicate population. Details in (see Methods)

**Fig. S4.**
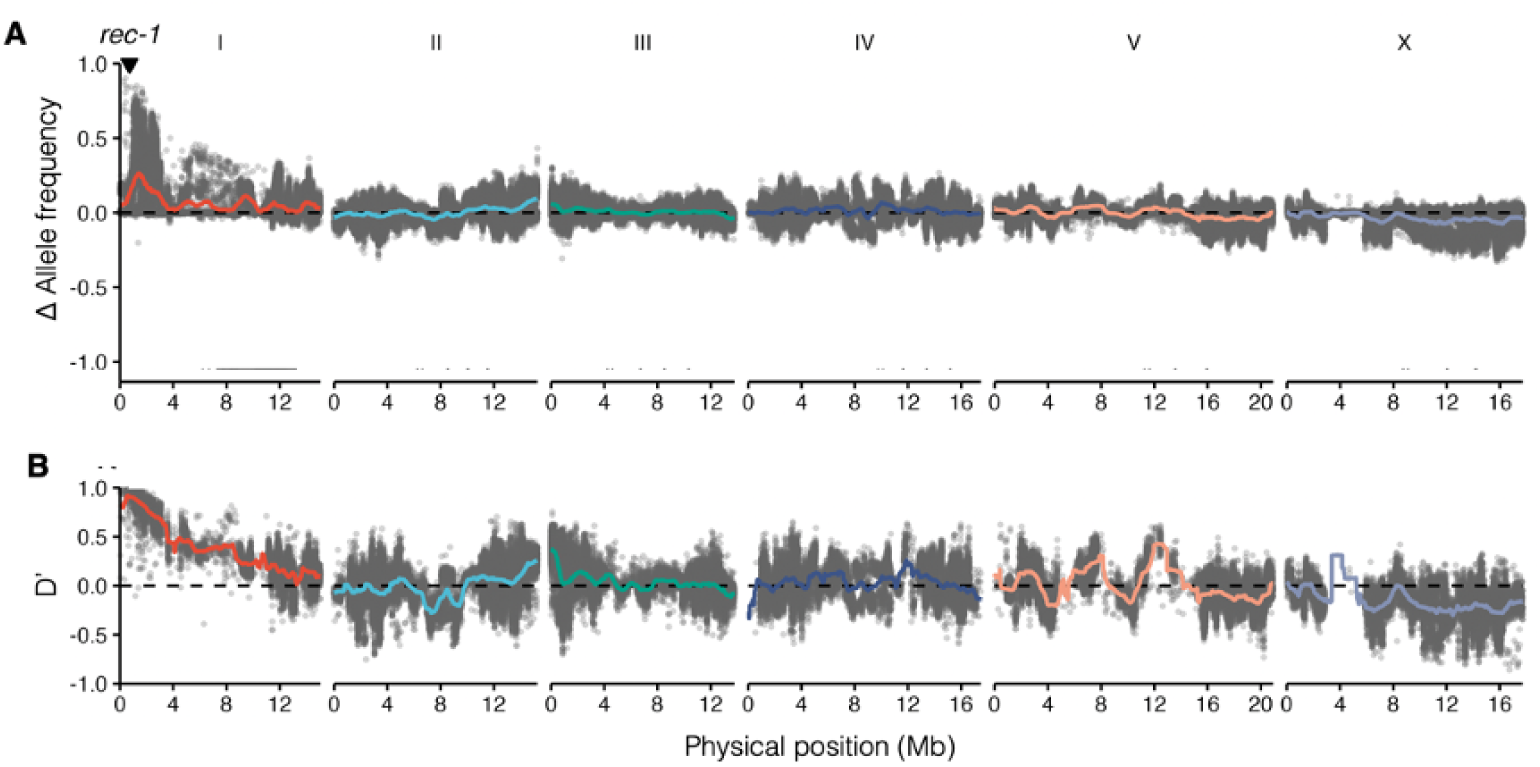
Ancestral difference between *rec-1* wild-type and mutant populations. **A.** *rec-1* sister populations with standing genetic variation were obtained through 340 crosses between the EEV1403 *rec-1* mutant strain and A6140 individuals (Figure S1, see Methods and Table S1 for strain and population designation and derivation). Points indicate differences in EEV1403 allele frequencies between the two sister populations. The *rec-1* locus is shown with an inverted triangle (I: 719,556 bp, WS245 genome). **B.** Linkage disequilibrium *D^′^* [63] between the *rec-1* mutant allele and the EEV1403 alleles among the wild-type and mutant ancestral sister populations (HR0 and MR0).

**Fig. S5.**
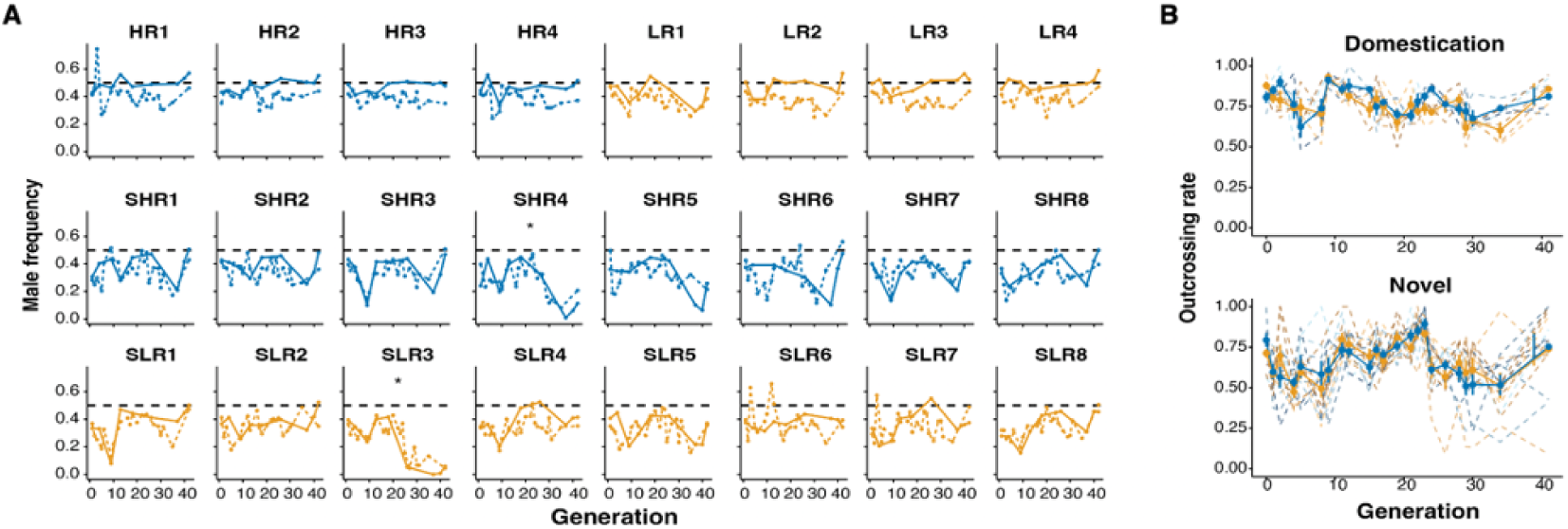
Outcrossing rates during the evolution of wild-type (blue) and mutant (orange) populations in experiment II. **A.** For each replicate population, male frequencies were obtained by visually scoring sexes (solid lines) or by videotaping worms and extracting phenotypes with the multi-worm tracker (dashed lines) during experimental evolution section 4. Populations that evolved in the domestication environment are shown in the first row, and those evolved in the novel salt environment in the second and third rows. SLR3 and SHR4 populations (asterisks) showed male frequencies compared to the average across all populations (Linear Mmodel: *outcrossing∼ population*+*generation*). The adaptation assays excluded these populations (Table S4). **B.**. The outcrossing rate (2*×* the male frequency in the subsequent generation [85], during experimental evolution in the domestication (upper panel) and novel environments (lower panel) for *rec-1* wild-type (blue) and mutant (red) populations (dashed = population; solid = average with standard error). Average outcrossing rates between the domestication and novel environments differ, with values of 0.66 and 0.77, respectively (Welch’s t-test: *p*-value = 9.8 *×* 10*^−^*^7^). No significant difference was found between the *rec-1* wild-type and mutant populations (Welch’s t-test).

**Fig. S6.**
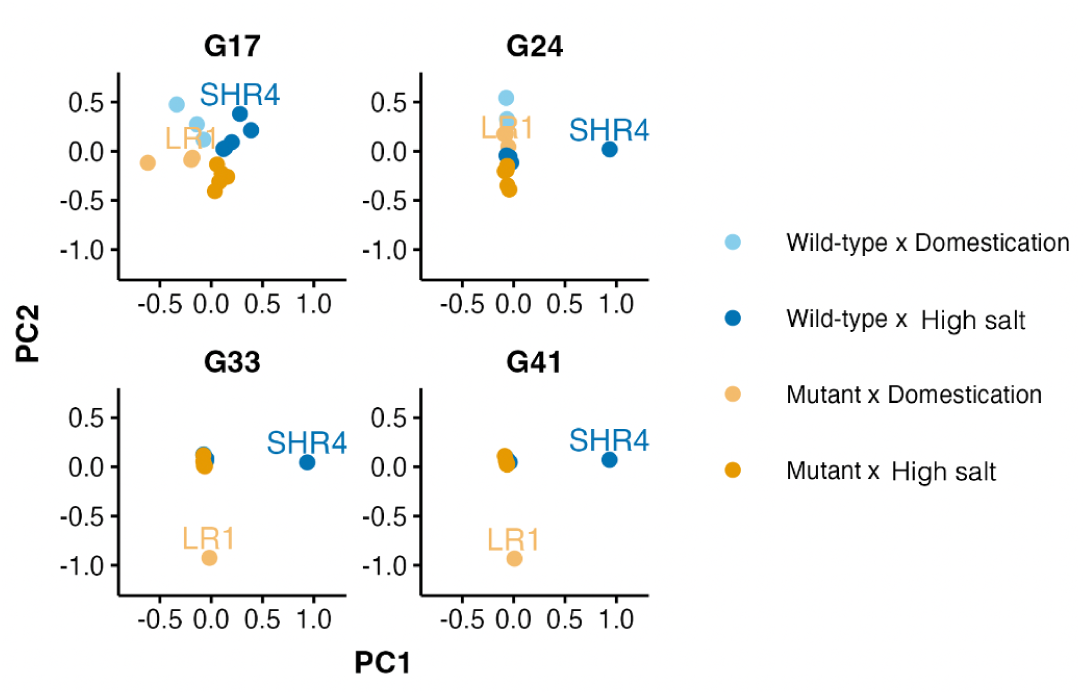
Principal component analysis of SNV allele frequency change. PCA was done on the allele frequency change from the first generation of each SNV in all sequenced populations Table S4. Allele frequencies were interpolated using the adjacent time points when samples were missing. Populations that evolved in the domestication (high-salt) environment are indicated by light (dark) colors and wild-type (mutant) populations in blue (orange). The SHR4 and LR1 populations appear as outliers in this analysis. During experimental evolution, the SHR4 population significantly decreased outcrossing rates (Figure S5), while the LR1 population faced the invasion of obligate self-fertilization mutants (Figure S7), and so were excluded from subsequent genomic analysis at the problematic time points (Table S4).

**Fig. S7.**
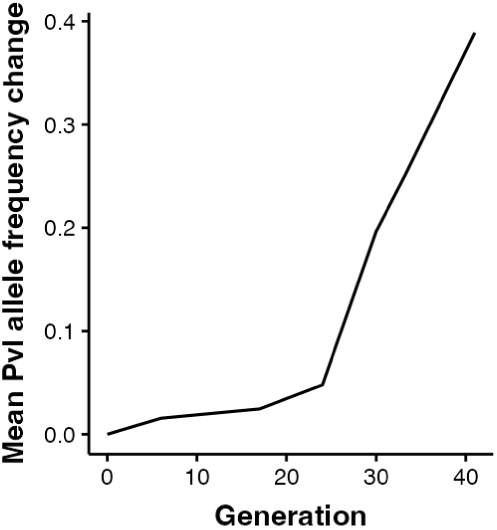
Protruding vulva phenotype. The protruding-vulva (Pvl) mutant phenotype [92] was observed in the LR1, LR2, and SLR3 populations. The Pvl phenotype results from a developmental defect in the vulva. Defects in the vulva can lead to a “bag-of-worms” phenotype, where the L1 larvae hatch within the mother. Additionally, individuals with this phenotype are obligate selfers, as they cannot outcross with males. The LR1 population, the only sequenced population among the three populations harboring individuals with the Pvl phenotype, is an outlier among the sequenced population after the 24th generation (Figure S6). This coincides with an increase of Pvl individuals in the LR1 population. Pvl individual proportion was estimated by sequencing PvL individuals after the 40th generation and measuring the average allele frequency of the initially rare PvL allele within the pool-sequenced LR1 samples (initial frequency in the LR1 population *<* 20%).

**Fig. S8.**
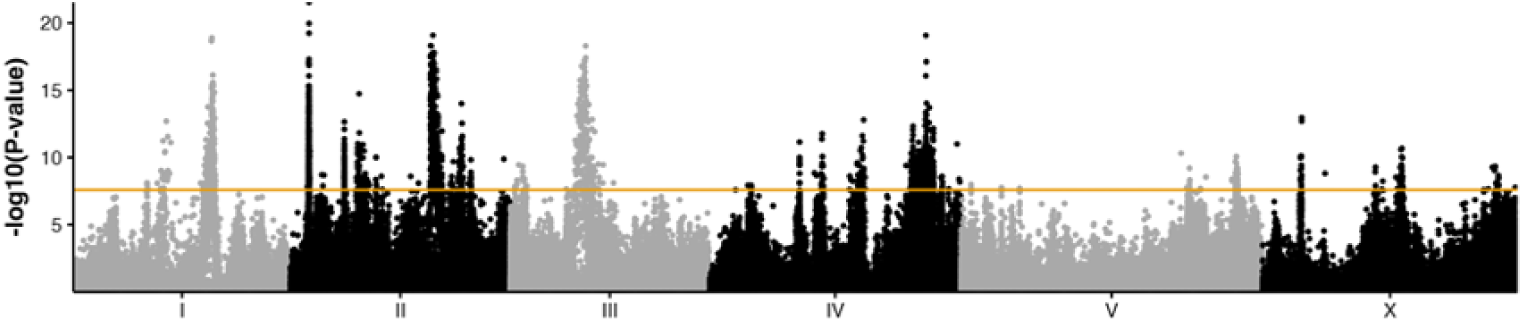
Effect of environment on SNV differentiation during experiment II. Environment specific-response, measured as the significance of consistent single-SNV allele frequency changes among replicate populations relative to generation zero (binomial generalized linear model, GLM allelic count *∼* generation * environment; Welch t-test on generation:environment interaction; see section 4). The orange line indicates the *α* = 0.05 threshold obtained by 1,000 random permutations of generation within each replicate population.

**Fig. S9.**
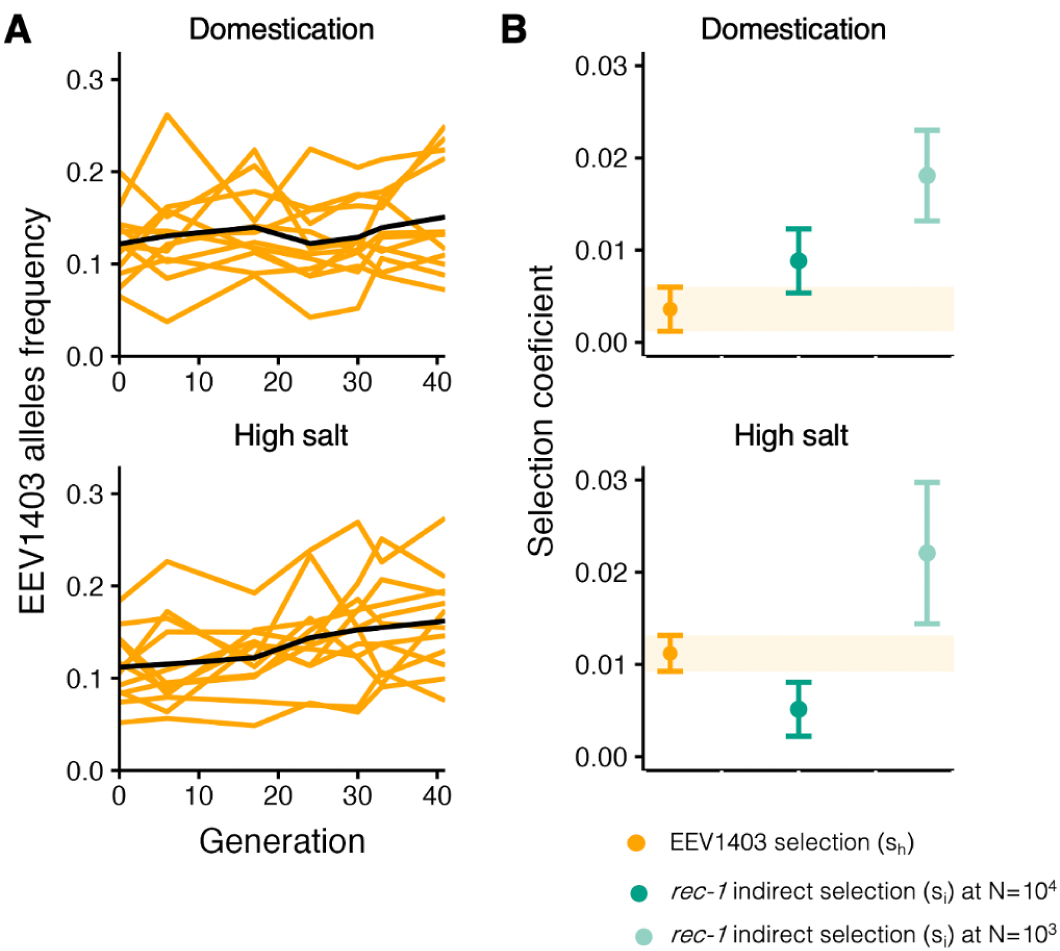
Selection of the EEV1403 haplotype is too weak to explain the indirect selection of *rec-1*. We asked that selection of the EEV1403 introgressed haplotype linked to *rec-1* (Figure S10) do not explain the selection at *rec-1*. **A.** To obtain an estimation of the selection of the EEV1403 haplotype without the *rec-1* mutant, we measured allele frequency changes of SNV marking the EEV1403 haplotype in the *rec-1* wild-type population (HR) used in experiment II. **B.** The selection coefficient (*s_h_*), of the EEV1403 haplotype (orange) is inferred from average allele frequency trajectories. Comparison of mean and 83% CI show that it is weaker than the indirect selection of the *rec-1* mutant (*s_i_*, green), smaller population sizes.

**Fig. S10.**
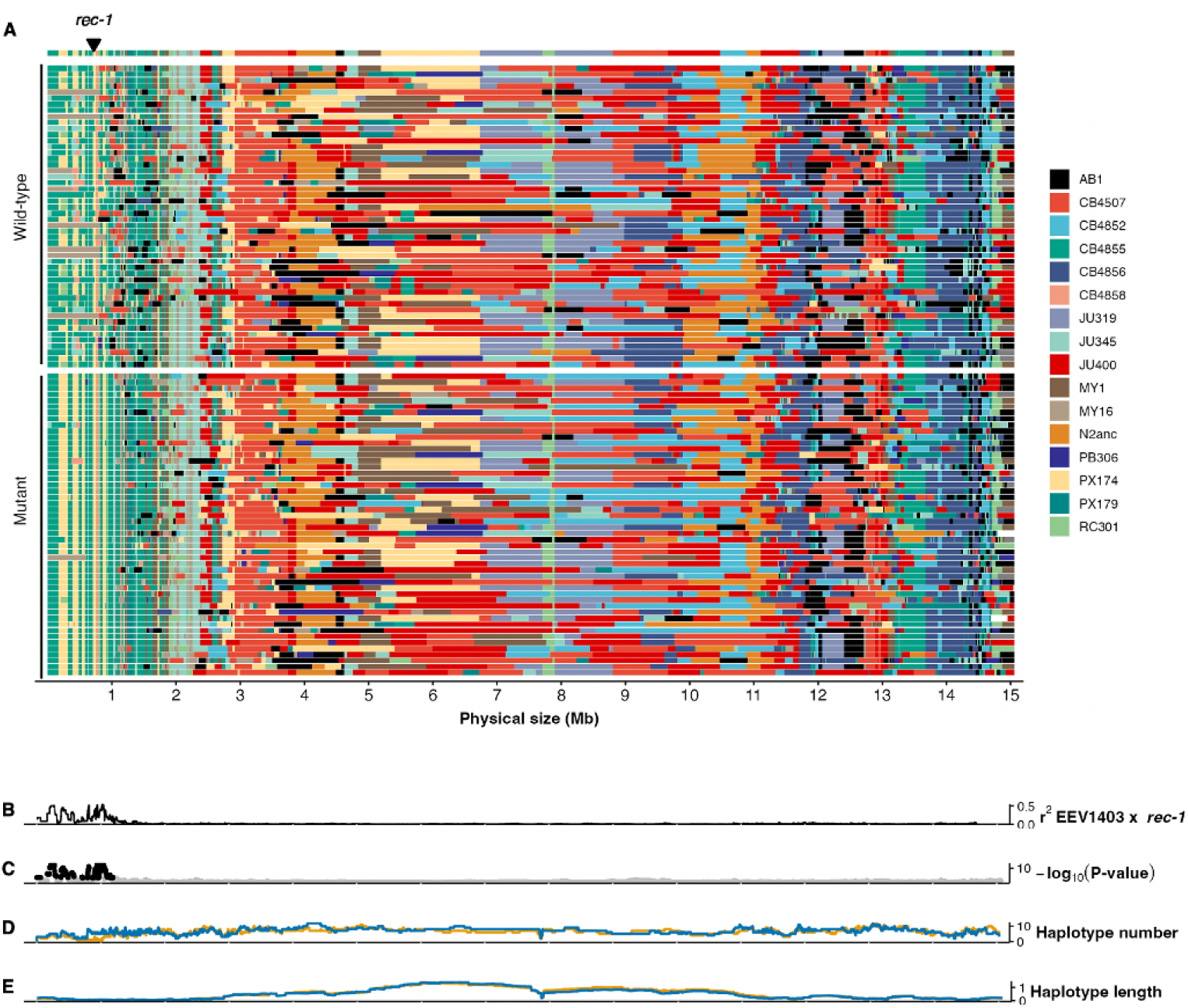
Haplotype founder blocks at chromosome I in the evolved *rec-1* polymorphic populations. Inferred founder genotypes (colors) in 50 wild-type and 50 mutant inbred lines (rows) derived from three *rec-1* polymorphic populations following evolution in the salt environment under the small population size treatment (Table S8). The EEV1403 strain, used to generate the *rec-1* wild-type and mutant populations, is shown as the top row (Figure 4a). **B.** Linkage disequilibrium (*r*^2^; squared correlation) between the *rec-1* locus and EEV1403 genotypes.**C.** Significance of the difference in genotype frequencies between *rec-1* wild-type and mutant lines (Linear Model: rec1 0 + haplotype; p-value obtained with F-test). Significant values are shown in black, and not significant values in grey (*α* = 0.05); with thresholds obtained by 1,000 random permutations of *rec-1* identity). **D.** Haplotype number of different founder genotypes in *rec-1* wild-type (blue) or mutant lines along physical distances (1kb intervals). **E.** Mean length (Mb) of haplotype extending 1kb in *rec-1* wild-type (blue) or mutant lines.

**Fig. S11.**
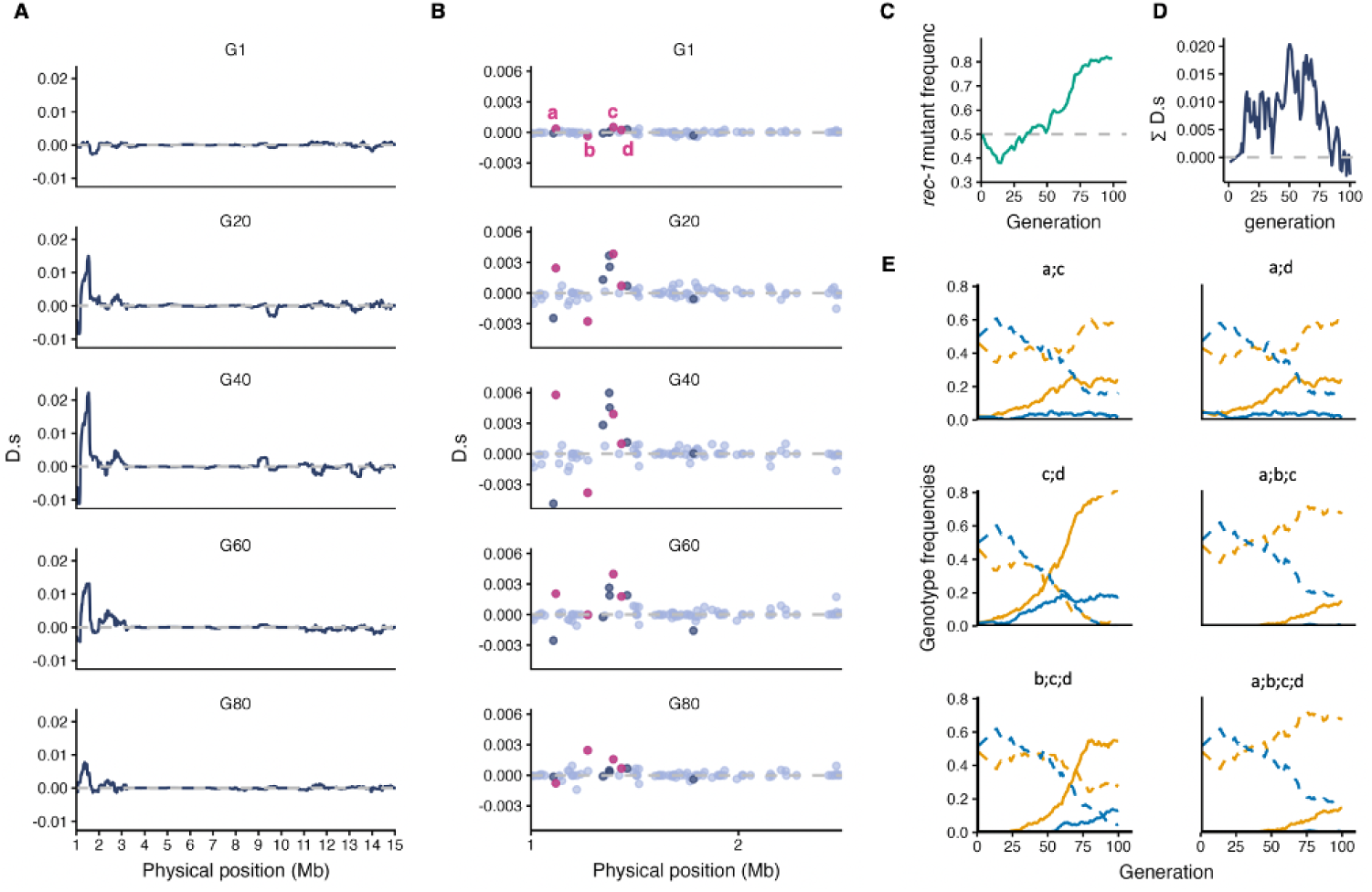
Linkage disequilibrium between *rec-1* and nearby genotypes. **A,B.** Association of *rec-1* and beneficial alleles on chromosome I during a single simulation run where the *rec-1* mutant is favored. Associations are highlighted by the *D.s*metric, the linkage disequilibrium (D) of beneficial with the *rec-1* mutant allele weighted by their selection coefficient (s). *D.s* is positive when the *rec-1* mutant is associated with beneficial alleles. **A.** *D.s* values are summed in a 1Mb sliding window and averaged over generations. **B.** the *D.s* value for each SNV over generations within the interval from 1 to 2.5Mb of chromosome I. SNVs with higher *D.s* are highlighted with a darker shade of purple or pink. The pink dots indicate a few SNV shown in panel *E*. **C.** Allele frequency of the *rec-1* mutant over generations. **D**. *D.s* summed genome-wide over generations. **E**. Frequency over a generation of two, three, and four beneficial allele combinations for the previously highlighted SNV (solid lines) and the alternative genotype combinations (dashed lines). Colors indicate whether the genotype is linked to the *rec-1* mutant (orange) or wild-type (blue) allele.

**Fig. S12.**
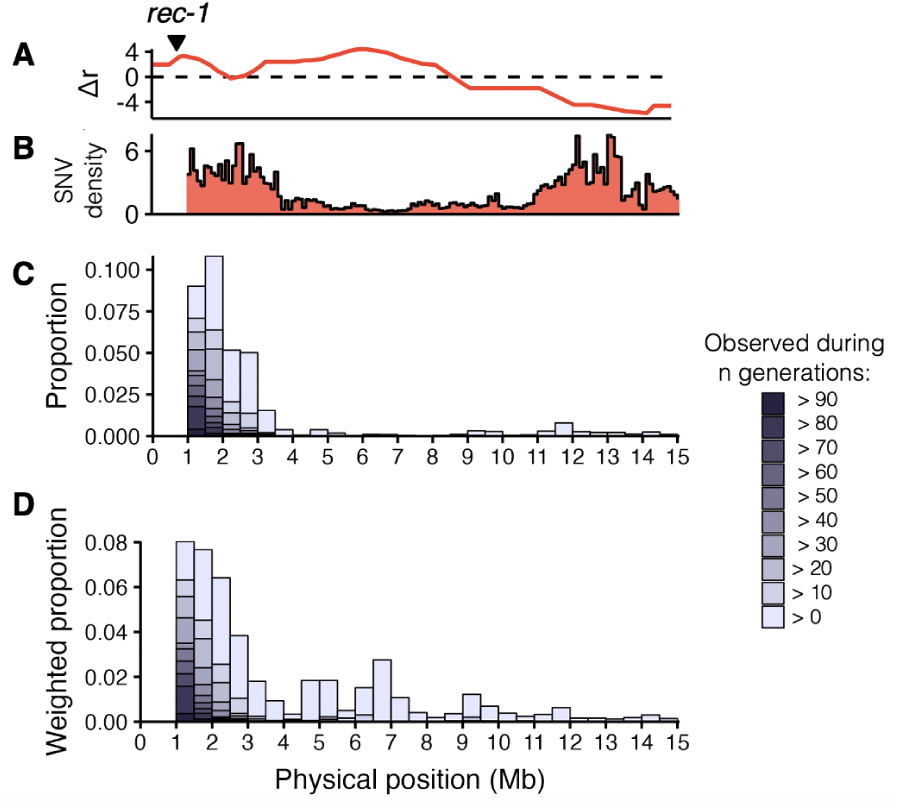
Physical distances between *rec-1* and favored genotypes. **A,B.** Δ*r* and SNV density from Figure 2 for chromosome I. The first Mb of the chromosome is depleted of genetic diversity in the simulations. **C.** Staked bars showing the proportion of SNV that are associated with the *rec-1* mutant across the genome (500 kb window size). An arbitrary linkage (*D.s*) threshold was set to determine which SNV is associated by taking the maximum observed value for chromosome II, which should not be associated with one of the *rec-1* alleles, found on chromosome I (Methods and Figure 4). The shade of purple indicates the number of generation during which the association is observed. **D.** SNV proportion associated with the *rec-1* mutant allele is inversely weighted by SNV density across the chromosome. Despite scarce genetic diversity, the *rec-1* mutant allele can be briefly associated with fitness loci in the central region of chromosome I.

**Fig. S13.**
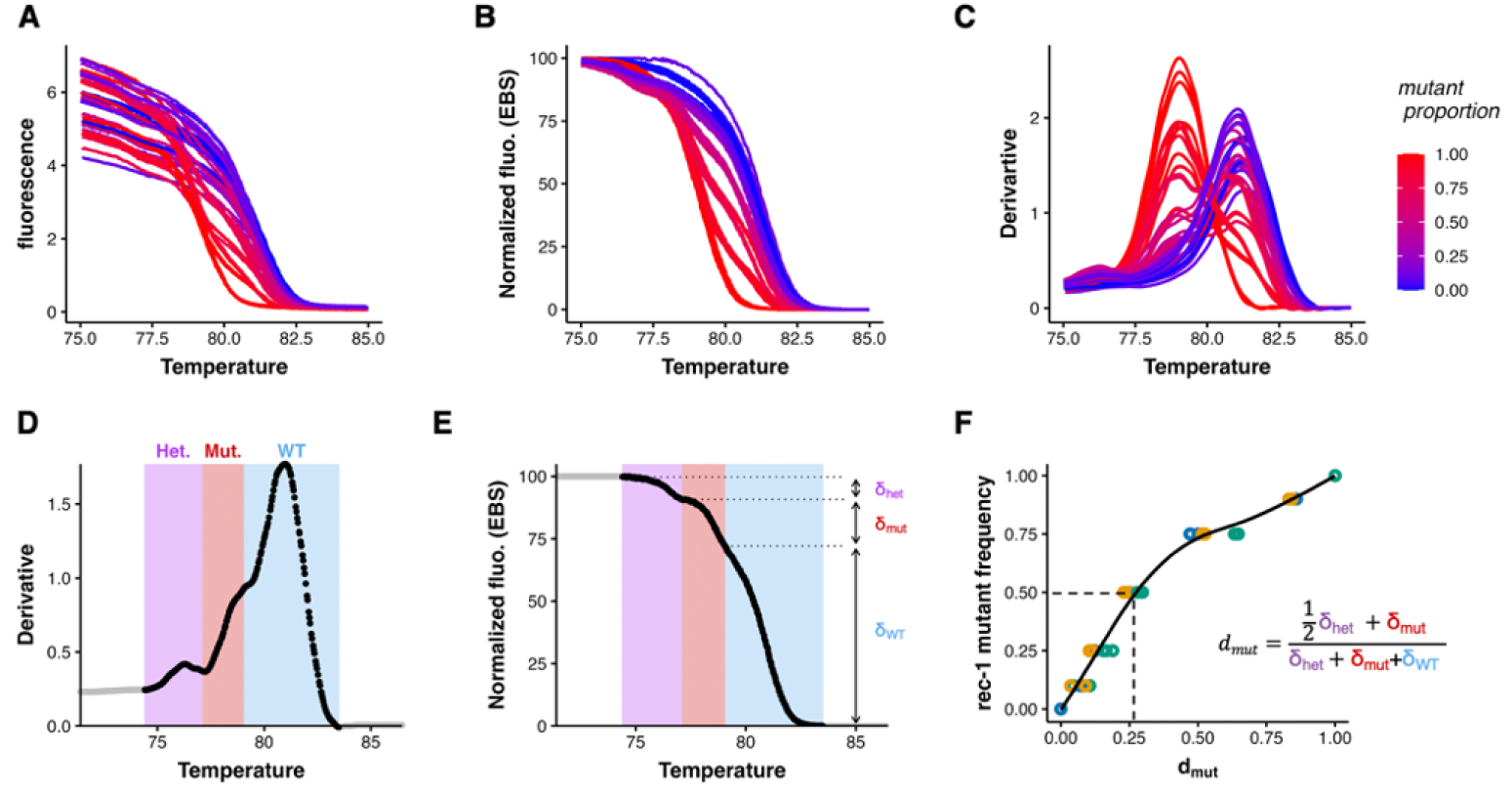
*rec-1* alleles quantification. **A.** The melting temperature (Tm) differs between the *rec-1* alleles, and different allelic proportions result in distinct melting curves. Disassociation between DNA molecules is measured by quantification of fluorescence during the qPCR section 4. **B.** Normalization of the melting curve to account for initial variability in fluorescence. **C.** The first derivative of normalized fluorescence on temperature. **D.** The first derivative for each sample is employed to detect different melting “phases” using a custom algorithm (Github) to delimit predominantly “heteroduplex” (purple), “mutant” (red), or “wild-type” (blue) phases. Phase decomposition is illustrated for a sample containing 1:1 mutant: wild-type individuals. **E.** Normalized fluorescence decay with temperature (*δ*) is quantified for each *rec-1* genotype pool. **F.** These metrics are used to calculate *dmut*, the fluorescence decay attributed to the mutant allele (equation within the figure). *dmut* is employed to predict the frequency of *rec-1* mutant alleles within samples using calibration samples. Different colors of the dots indicate three independent sets of calibration samples. The black lines show estimates from a generalized additive model of mutant allele frequency on *dmut* following ref. [93].

**Fig. S14.**
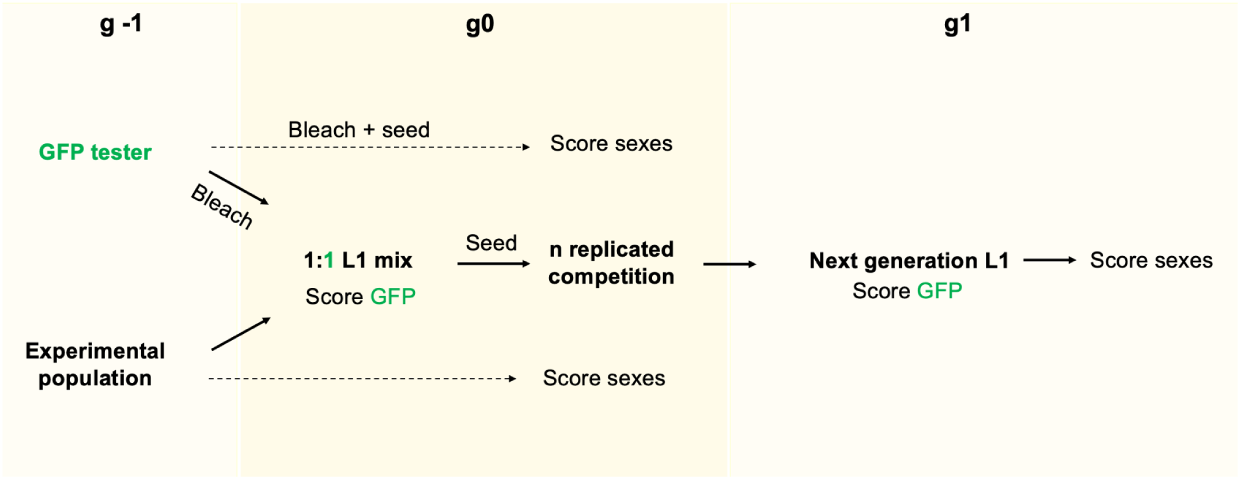
Adaptation assay design. Fitness of experimental populations is determined by competing them against the EEV1402 tester strain, a strain homozygous for a Green Fluorescent Protein (GFP) construct (see Methods). The experimental population and the tester undergo the bleach/hatch-off protocol (g-1), and recovered first larvae staged (L1) at generation zero (g0) are mixed in equal proportions. This mix is seeded into technical replicate plates, each with a population size of *N* = 3 *×* 10^3^. Fitness of the experimental population is quantified by assessing the initial proportion of GFP L1s in the initial mix (g0) and after one generation of competition (g1). To estimate GFP allele frequency at g1, the number of GFP heterozygotes at g1 is estimated considering the g0 and g1 sex ratio and the expected self-fertilization and outcrossing rates of the tester and experimental in Supplementary Table S7.

**Table S1.**
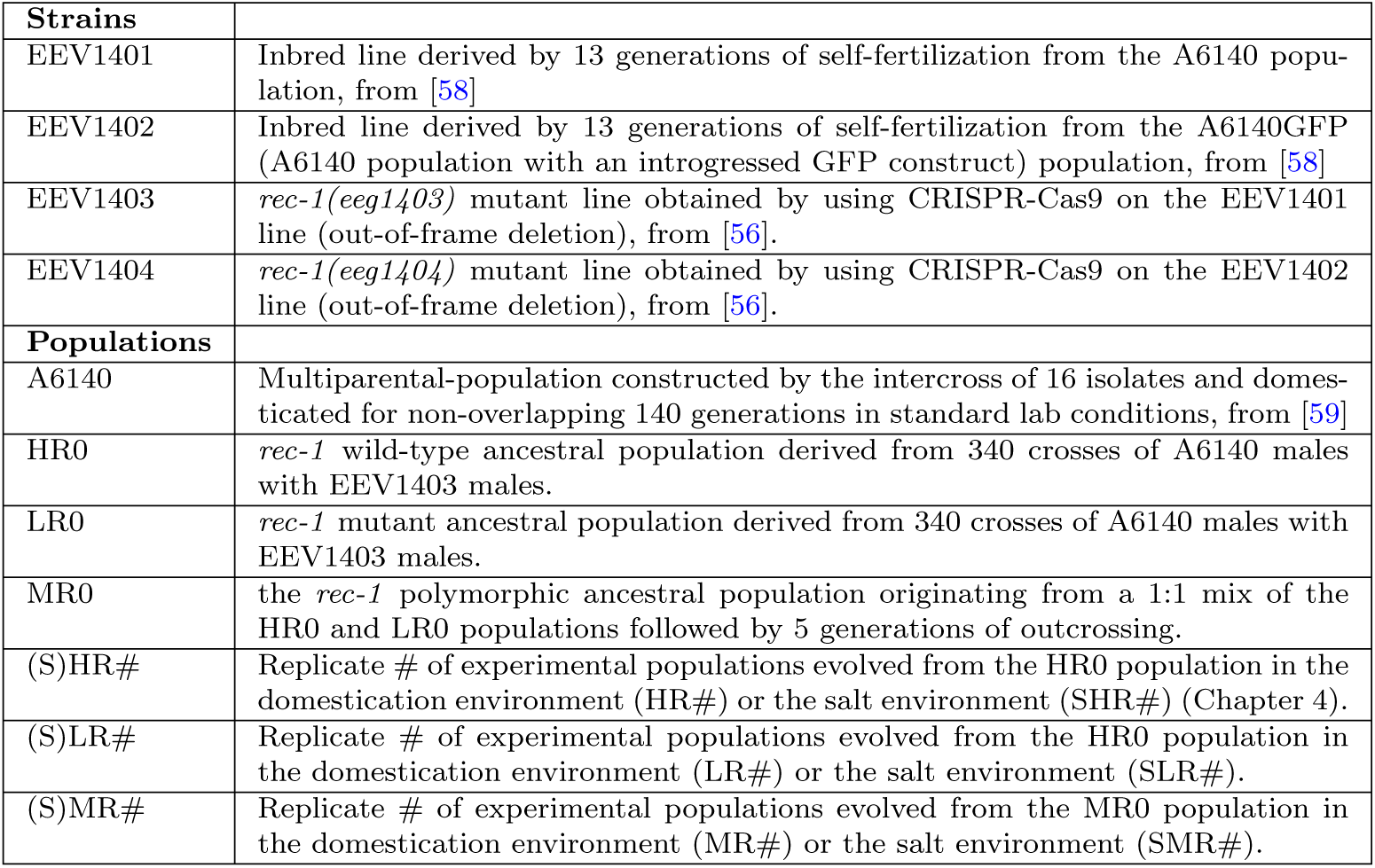
Strains and experimental population designation and description.

| **Strains** |  |
| --- | --- |
| EEV1401 | Inbred line derived by 13 generations of self-fertilization from the A6140 population, from [58] |
| EEV1402 | Inbred line derived by 13 generations of self-fertilization from the A6140GFP (A6140 population with an introgressed GFP construct) population, from [58] |
| EEV1403 | *rec-1(eeg1403)* mutant line obtained by using CRISPR-Cas9 on the EEV1401 line (out-of-frame deletion), from [56]. |
| EEV1404 | *rec-1(eeg1404)* mutant line obtained by using CRISPR-Cas9 on the EEV1402 line (out-of-frame deletion), from [56]. |
| **Populations** |  |
| A6140 | Multiparental-population constructed by the intercross of 16 isolates and domesticated for non-overlapping 140 generations in standard lab conditions, from [59] |
| HR0 | *rec-1* wild-type ancestral population derived from 340 crosses of A6140 males with EEV1403 males. |
| LR0 | *rec-1* mutant ancestral population derived from 340 crosses of A6140 males with EEV1403 males. |
| MR0 | the *rec-1* polymorphic ancestral population originating from a 1:1 mix of the HR0 and LR0 populations followed by 5 generations of outcrossing. |
| (S)HR# | Replicate # of experimental populations evolved from the HR0 population in the domestication environment (HR#) or the salt environment (SHR#) (Chapter 4). |
| (S)LR# | Replicate # of experimental populations evolved from the LR0 population in the domestication environment (LR#) or the salt environment (SLR#). |
| (S)MR# | Replicate # of experimental populations evolved from the MR0 population in the domestication environment (MR#) or the salt environment (SMR#). |

**Table S2.** Data and scripts availability.

| Content | Github path |
| --- | --- |
| Genotype Tables | rec1EE/genotype |
| Recombination rate differences ($\Delta r$) | rec1EE/recombinationMaps |
| *rec-1* mutant frequency measurement (melting curve analysis) | PAREE/melting_curves_analysis |
| Genomic heritability estimation | rec1EE/heritability |
| Haplotyping of inbred lines | rec1EE/haplotype |
| Genome scan for SNV differentiations result | rec1EE/candidateSelection |
| Fitness assay data and analysis | rec1EE/adaptation_fitness |
| Individual-based simulations (SLiM) | PAREE/SLiM |
| Male frequency estimation by Multiwormtracker | multiwormtracker/male |
| Outcrossing rate data | rec1EE/outcrossing&outliers |
| Allele frequency correlation | rec1EE/observed_r2 |
| Expected $r^2$ | rec1EE/predicted_r2 |
| Direct selection | rec1EE/direct_selection |
| Indirect selection | rec1EE/indirect_selection |
| Selection of the EEV1403 haplotype ($s_h$) | rec1EE/indirect_selection |

**Table S3.** Design of experiment I. testing for direct selection at *rec-1*.

| Environment | Background | WT strain | Mut. strain | Male freq. | Initial mut. freq. | Replicates |
| --- | --- | --- | --- | --- | --- | --- |
| Domestication | EEV1401 | EEV1401 | EEV1403 | high | 0.8 | 15 |
| Domestication | EEV1401 | EEV1401 | EEV1403 | low | 0.5 | 21 |
| Domestication | EEV1401 | EEV1401 | EEV1403 | low | 0.8 | 13 |
| Domestication | EEV1402 | EEV1402 | EEV1404 | high | 0.8 | 21 |
| Domestication | EEV1402 | EEV1402 | EEV1404 | low | 0.5 | 18 |
| Domestication | EEV1402 | EEV1402 | EEV1404 | low | 0.8 | 17 |
| salt | EEV1401 | EEV1401 | EEV1403 | low | 0.5 | 19 |
| salt | EEV1402 | EEV1402 | EEV1404 | low | 0.5 | 19 |
|  |  |  |  |  |  | 143 |

**Table S4.**
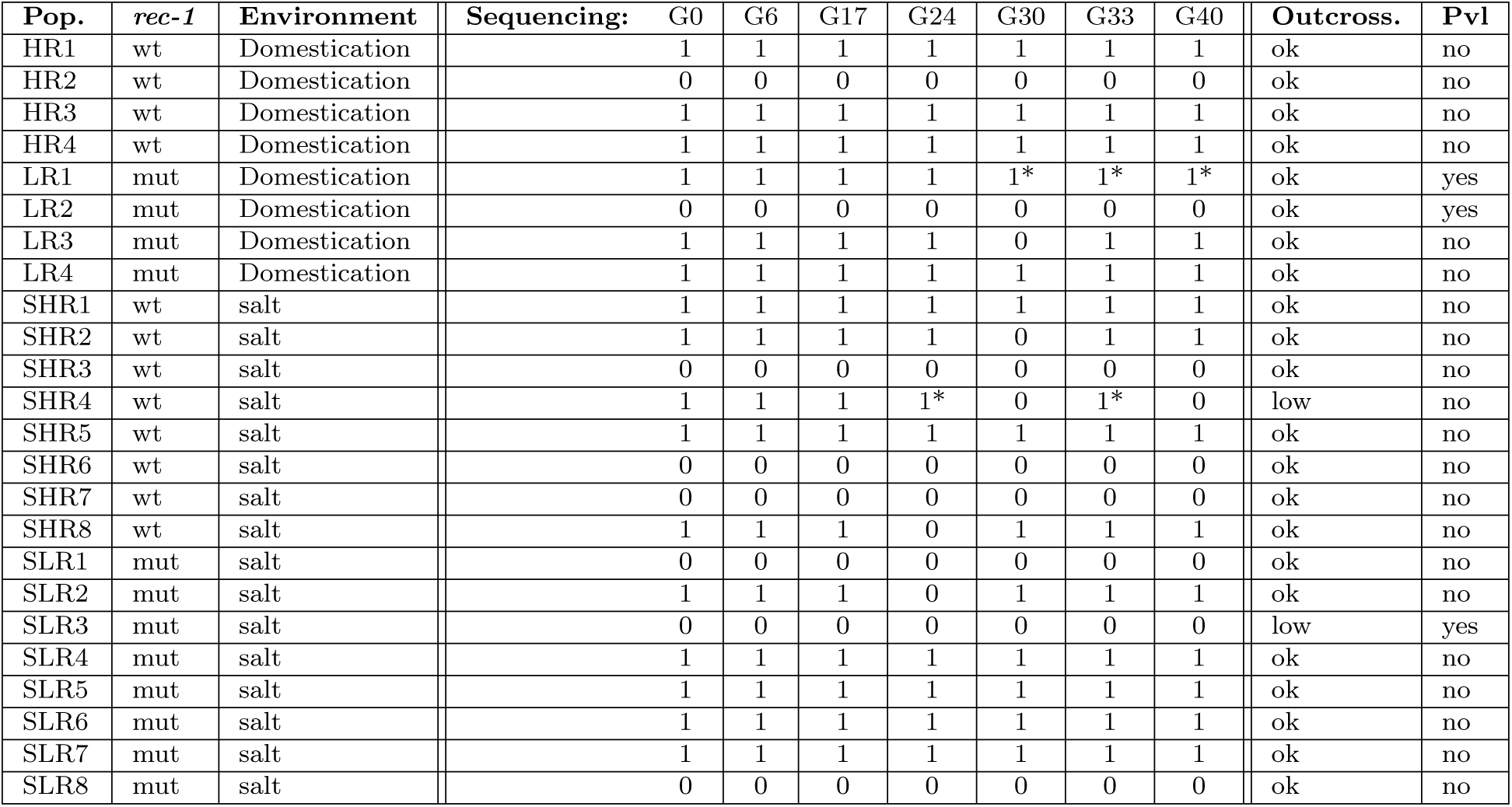
Experiment II. Columns show population designation and identity, as well as the time points during the experiment where individually pooled samples for whole-genome re-sequencing were obtained (indicated by a 1). The asterisks indicate outlier samples discarded from analysis (see Figure S6). Whether the population maintained high outcrossing (ok) or underwent a strong decrease in male frequency (low) is indicated (see Figure S5). Populations in which the Protruding-vulva phenotype (Pvl) was observed are also indicated Figure S7. All populations were tested for adaptation.

**Table S5.** Number of SNV in the pruned dataset to estimate genomic heritability for self-fertility.

| $\Delta r$ | chromosome | SNV number |
| --- | --- | --- |
| negative | I | 1911 |
| positive | I | 2367 |
| negative | II | 6699 |
| positive | II | 2036 |
| negative | III | 6986 |
| positive | III | 1224 |
| negative | IV | 5416 |
| positive | IV | 2699 |
| negative | V | 3350 |
| positive | V | 4683 |
| negative | X | 2392 |
| positive | X | 1016 |

**Table S6.** Number of technical replicates for the adaptation assays. Each experimental block (A,B,C,D) corresponds to an independent thaw and assay calendar date.

| Pop: | A | B | C | D |
| --- | --- | --- | --- | --- |
| HR0 | 5 | 10 | 3 | 3 |
| HR1 | 5 | 5 | 3 | 3 |
| HR2 | 5 | 5 | 3 | 3 |
| HR3 | 5 | 5 | 3 | 3 |
| HR4 | 5 | 5 | 3 | 3 |
| LR0 | 5 | 10 | 3 | 3 |
| LR1 | 5 | 5 | 3 | 3 |
| LR2 | 5 | 5 | 3 | 3 |
| LR3 | 5 | 5 | 3 | 3 |
| LR4 | 5 | 5 | 3 | 3 |
| SHR0 | 5 | 10 | 3 | 3 |
| SHR1 | 5 | 5 | 3 | 3 |
| SHR2 | 5 | 5 | 3 | 3 |
| SHR3 | 5 | 5 | 3 | 3 |
| SHR5 | 5 | 5 | 3 | 3 |
| SHR6 | 5 | 5 | 3 | 3 |
| SHR7 | 5 | 5 | 3 | 3 |
| SHR8 | 0 | 5 | 3 | 3 |
| SLR0 | 5 | 10 | 3 | 3 |
| SLR1 | 5 | 5 | 3 | 3 |
| SLR2 | 5 | 5 | 3 | 3 |
| SLR4 | 5 | 5 | 3 | 0 |
| SLR5 | 5 | 5 | 3 | 3 |
| SLR6 | 5 | 5 | 3 | 3 |
| SLR7 | 5 | 5 | 3 | 3 |
| SLR8 | 0 | 5 | 3 | 3 |

**Table S7.** Adaptation assays mating table. The expression in each cell corresponds to the expected proportion of progeny from the corresponding parents. The first line corresponds to the outcrossed progeny. It depends on each parent’s fitness (*W*) and proportion (*p*0), as well as the outcrossing rate (1 *− S*) calculated as twice the number of males [85]. The second line in cells [1,1] and [2,2] corresponds to the number of selfed progeny. These numbers depend only on the parental hermaphrodite fitness and the self-fertilization rate (*F*). Cells [1,2] and [2,1] are the green fluorescent protein (GFP) heterozygous progeny. Cell [1,1] is the GFP homozygous progeny. Cell [2,2] is the wild-type (WT) progeny. As GFP is dominant, only the proportion of WT progeny equals the proportion of scored WT L1 after the competition. This equivalence allows us to resolve the *W_W T_* term and the *W_GF P_* term, quantifying the proportion of GFP heterozygous individuals.

|  | **GFP herm** | **WT herm** |
| --- | --- | --- |
| **GFP male**<br>or self-fertilization | $p0_{GFP.male} \times p0_{GFP.herm} \times W_{GFP}^2 \times (1 - S)$<br>$+ F \times p0_{GFP.herm} \times W_{GFP}$ | $p0_{GFP.male} \times p0_{WT.herm} \times W_{GFP} \times W_{WT} \times (1 - S)$ |
| **WT male**<br>or self-fertilization | $p0_{WT.male} \times p0_{GFP.herm} \times W_{GFP} \times W_{WT} \times (1 - S)$ | $p0_{WT.male} \times p0_{WT.herm} \times W_{WT}^2 \times (1 - S)$<br>$+ S \times p0_{WT.herm} \times W_{WT}$ |

**Table S8.** Design of experiment III testing for indirect selection at *rec-1*. Inbred lines were derived from the three populations highlighted with an asterisk.

| Population | Environment | Census pop. size | Block | n. generation |
| --- | --- | --- | --- | --- |
| MR1 | Domestication | 10000 | 1 | 16 |
| MR3 | Domestication | 10000 | 1 | 16 |
| MR5 | Domestication | 10000 | 1 | 16 |
| MR7 | Domestication | 10000 | 2 | 14 |
| MR8 | Domestication | 10000 | 2 | 14 |
| MR9 | Domestication | 10000 | 2 | 14 |
| MR10 | Domestication | 10000 | 2 | 14 |
| MR11 | Domestication | 1000 | 2 | 14 |
| MR12 | Domestication | 1000 | 2 | 14 |
| MR13 | Domestication | 1000 | 2 | 14 |
| MR14 | Domestication | 1000 | 2 | 14 |
| MR15 | Domestication | 1000 | 2 | 14 |
| MR16 | Domestication | 1000 | 2 | 14 |
| MR17 | Domestication | 1000 | 2 | 14 |
| MR18 | Domestication | 1000 | 2 | 14 |
| MR19 | Domestication | 1000 | 2 | 14 |
| MR20 | Domestication | 1000 | 2 | 14 |
| MR21 | Domestication | 1000 | 2 | 14 |
| MR22 | Domestication | 1000 | 2 | 14 |
| MR23 | Domestication | 1000 | 2 | 14 |
| MR24 | Domestication | 1000 | 2 | 14 |
| MR25 | Domestication | 1000 | 2 | 14 |
| MR26 | Domestication | 1000 | 2 | 14 |
| MR27 | Domestication | 1000 | 2 | 14 |
| MR28 | Domestication | 1000 | 2 | 14 |
| MR29 | Domestication | 1000 | 2 | 14 |
| MR30 | Domestication | 1000 | 2 | 14 |
| SMR1 | salt | 10000 | 1 | 16 |
| SMR3 | salt | 10000 | 1 | 16 |
| SMR5 | salt | 10000 | 1 | 16 |
| SMR7 | salt | 10000 | 1 | 16 |
| SMR9 | salt | 10000 | 1 | 16 |
| SMR11 | salt | 10000 | 1 | 16 |
| SMR13 | salt | 10000 | 1 | 16 |
| SMR15 | salt | 10000 | 1 | 16 |
| SMR17 | salt | 10000 | 1 | 16 |
| SMR19 | salt | 10000 | 2 | 14 |
| SMR20 | salt | 10000 | 2 | 14 |
| SMR21 | salt | 10000 | 2 | 14 |
| SMR22 | salt | 10000 | 2 | 14 |
| SMR23 | salt | 1000 | 2 | 14 |
| SMR24* | salt | 1000 | 2 | 14 |
| SMR25* | salt | 1000 | 2 | 14 |
| SMR26 | salt | 1000 | 2 | 14 |
| SMR27 | salt | 1000 | 2 | 14 |
| SMR28 | salt | 1000 | 2 | 14 |
| SMR29 | salt | 1000 | 2 | 14 |
| SMR30 | salt | 1000 | 2 | 14 |
| SMR31 | salt | 1000 | 2 | 14 |
| SMR32 | salt | 1000 | 2 | 14 |
| SMR33 | salt | 1000 | 2 | 14 |
| SMR34 | salt | 1000 | 2 | 14 |
| SMR35 | salt | 1000 | 2 | 14 |
| SMR36 | salt | 1000 | 2 | 14 |
| SMR37 | salt | 1000 | 2 | 14 |
| SMR38 | salt | 1000 | 2 | 14 |
| SMR39 | salt | 1000 | 2 | 14 |
| SMR40* | salt | 1000 | 2 | 14 |
| SMR41 | salt | 1000 | 2 | 14 |
| SMR42 | salt | 1000 | 2 | 14 |

## References

[1] Lenormand T, Engelstädter J, Johnston SE, Wijnker E, Haag CR. Evolutionary mysteries in meiosis. Philosophical Transactions of the Royal Society B: Biological Sciences. 2016;371(1706):20160001.

[2] Stapley J, Feulner PG, Johnston SE, Santure AW, Smadja CM. Variation in recombination frequency and distribution across eukaryotes: patterns and processes. Philosophical Transactions of the Royal Society B: Biological Sciences. 2017;372(1736):20160455.

[3] Baker Z, Schumer M, Haba Y, Bashkirova L, Holland C, Rosenthal GG, et al. Repeated losses of PRDM9-directed recombination despite the conservation of PRDM9 across vertebrates. Elife. 2017;6:e24133.

[4] Brand CL, Cattani MV, Kingan SB, Landeen EL, Presgraves DC. Molecular evolution at a meiosis gene mediates species differences in the rate and patterning of recombination. Current Biology. 2018;28(8):1289–1295.

[5] Comeron JM, Ratnappan R, Bailin S. The many landscapes of recombination in Drosophila melanogaster. PLoS Genet. 2012;8(10):e1002905.

[6] Zhang L, Liang Z, Hutchinson J, Kleckner N. Crossover patterning by the beamfilm model: analysis and implications. PLoS genetics. 2014;10(1):e1004042.

[7] Haenel Q, Laurentino TG, Roesti M, Berner D. Meta-analysis of chromosomescale crossover rate variation in eukaryotes and its significance to evolutionary genomics. Molecular ecology. 2018;27(11):2477–2497.

[8] Brazier T, Glémin S. Diversity and determinants of recombination landscapes in flowering plants. PLoS Genetics. 2022;18(8):e1010141.

[9] Sardell JM, Kirkpatrick M. Sex differences in the recombination landscape. The American Naturalist. 2020;195(2):361–379.

[10] Woodruff GC, Teterina AA. Degradation of the Repetitive Genomic Landscape in a Close Relative of Caenorhabditis elegans. Molecular Biology and Evolution. 2020;37(9):2549–2567.

[11] Nam K, Ellegren H. Recombination drives vertebrate genome contraction. PLoS genetics. 2012;8(5):e1002680.

[12] Gerstein MB, Lu ZJ, Van Nostrand EL, Cheng C, Arshinoff BI, Liu T, et al. Integrative analysis of the Caenorhabditis elegans genome by the modENCODE project. Science. 2010;330(6012):1775–1787.

[13] Kent TV, Uzunović J, Wright SI. Coevolution between transposable elements and recombination. Philosophical Transactions of the Royal Society B: Biological Sciences. 2017;372(1736):20160458.

[14] Wolf JB, Ellegren H. Making sense of genomic islands of differentiation in light of speciation. Nature Reviews Genetics. 2017;18(2):87–100.

[15] Cutter AD, Payseur BA. Genomic signatures of selection at linked sites: unifying the disparity among species. Nature Reviews Genetics. 2013;14(4):262–274.

[16] Begun DJ, Aquadro CF. Levels of naturally occurring DNA polymorphism correlate with recombination rates in D. melanogaster. Nature. 1992;356(6369):519– 520.

[17] Hunt P. Meiosis in mammals: recombination, non-disjunction and the environment. Biochemical Society Transactions. 2006;34(4):574–577.

[18] Hassold T, Hunt P. To err (meiotically) is human: the genesis of human aneuploidy. Nature Reviews Genetics. 2001;2(4):280–291.

[19] Fernandes JB, Séguéla-Arnaud M, Larchev^eque C, Lloyd AH, Mercier R. Unleashing meiotic crossovers in hybrid plants. Proceedings of the National Academy of Sciences. 2018;115(10):2431–2436.

[20] Fisher RA.: The genetical theory of natural selection. Clarendon. Oxford.

[21] Barton NH. A general model for the evolution of recombination. Genetics Research. 1995;65(2):123–144.

[22] Burt A. Perspective: Sex, recombination and the efficacy of selection: was Weissman right? Evolution. 2000;54:337–351.

[23] Barton N. Why sex and recombination? Cold Spring Harbor Symposia on Quantitative Biology. 2009;74:187–195.

[24] Roze D. A simple expression for the strength of selection on recombination generated by interference among mutations. Proceedings of the National Academy of Sciences. 2021;118(19):e2022805118.

[25] Otto SP, Lenormand T. Resolving the paradox of sex and recombination. Nat Rev Genet. 2002;3(4):252–61.

[26] Eshel I, Feldman MW. On the evolutionary effect of recombination. Theor Popul Biol. 1970;1(1):88–100.

[27] Neher RA, Shraiman BI. Competition between recombination and epistasis can cause a transition from allele to genotype selection. Proc Natl Acad Sci U S A. 2009;106(16):6866–71.

[28] Kouyos RD, Silander OK, Bonhoeffer S. Epistasis between deleterious mutations and the evolution of recombination. Trends in ecology & evolution. 2007;22(6):308–315.

[29] Hill WG, Robertson A. The effect of linkage on limits to artificial selection. Genetics Research. 1966;8(3):269–294.

[30] Barton NH, Otto SP. Evolution of recombination due to random drift. Genetics. 2005;169(4):2353–70.

[31] Otto SP. Selective interference and the evolution of sex. Journal of Heredity. 2021;112(1):9–18.

[32] Roze D, Barton NH. The Hill-Robertson effect and the evolution of recombination. Genetics. 2006;173(3):1793–811.

[33] Charlesworth B, Barton NH. Recombination load associated with selection for increased recombination. Genet Res Camb. 1996;67(1):27–41.

[34] Rice WR, Chippindale AK. Sexual recombination and the power of natural selection. Science. 2001;294(5542):555–9.

[35] Goddard MR, Godfray HC, Burt A. Sex increases the efficacy of natural selection in experimental yeast populations. Nature. 2005;434(7033):636–640.

[36] McDonald MJ, Rice DP, Desai MM. Sex speeds adaptation by altering the dynamics of molecular evolution. Nature. 2016;531(7593):233–236.

[37] Weissman DB, Barton N. Limits to the rate of adaptive substitution in sexual populations. PLoS Genet. 2012;8:e1002740.

[38] Bernstein H, Bernstein C, Michod RE. Meiosis as an evolutionary adaptation for DNA repair. DNA repair. 2011;11:357.

[39] Kosheleva K, Desai MM. Recombination alters the dynamics of adaptation on standing variation in laboratory yeast populations. Molecular Biology and Evolution. 2018;35(1):180–201.

[40] Becks L, Agrawal AF. The evolution of sex is favored during adaptation to new environments. PLoS Biol. 2012;10(5):e1001317.

[41] Weissman DB, Hallatschek O. The rate of adaptation in large sexual populations with linear chromosomes. Genetics. 2014;196(4):1167–1183.

[42] Barghi N, Hermisson J, Schlotterer C. Polygenic adaptation: a unifying framework to understand positive selection. Nature Reviews Genetics. 2020;.

[43] Barrett RD, Schluter D. Adaptation from standing genetic variation. Trends in ecology & evolution. 2008;23(1):38–44.

[44] Otto SP, Barton NH. The evolution of recombination: removing the limits to natural selection. Genetics. 1997;147(2):879–906.

[45] Rockman MV, Kruglyak L. Recombinational landscape and population genomics of Caenorhabditis elegans. PLoS genetics. 2009;5(3):e1000419.

[46] Noble LM, Yuen J, Stevens L, Moya N, Persaud R, Moscatelli M, et al. Selfing is the safest sex for Caenorhabditis tropicalis. Elife. 2021;10:e62587.

[47] Ross JA, Koboldt DC, Staisch JE, Chamberlin HM, Gupta BP, Miller RD, et al. Caenorhabditis briggsae recombinant inbred line genotypes reveal interstrain incompatibility and the evolution of recombination. PLoS genetics. 2011;7(7):e1002174.

[48] Teterina AA, Willis JH, Lukac M, Jovelin R, Cutter AD, Phillips PC. Genetic diversity landscapes in outcrossing and selfing Caenorhabditis nematodes. bioRxiv. 2022;p. 2022–12.

[49] Crombie TA, Zdraljevic S, Cook DE, Tanny RE, Brady SC, Wang Y, et al. Deep sampling of Hawaiian Caenorhabditis elegans reveals high genetic diversity and admixture with global populations. Elife. 2019;8.

[50] Andersen EC, Gerke JP, Shapiro JA, Crissman JR, Ghosh R, Bloom JS, et al. Chromosome-scale selective sweeps shape Caenorhabditis elegans genomic diversity. Nat Genet. 2012;44(3):285–90.

[51] Rockman MV, Skrovanek SS, Kruglyak L. Selection at linked sites shapes heritable phenotypic variation in C. elegans. Science. 2010;330(6002):372–6.

[52] Yoshida K, Rodelsperger C, Roseler W, Riebesell M, Sun S, Kikuchi T, et al. Chromosome fusions repatterned recombination rate and facilitated reproductive isolation during Pristionchus nematode speciation [Journal Article]. Nat Ecol Evol. 2023;7(3):424–439.

[53] Rose A, Baillie D. A mutation in Caenorhabditis elegans that increases recombination frequency more than threefold. Nature. 1979;281(5732):599–600.

[54] Zetka MC, Rose AM. Mutant rec-1 eliminates the meiotic pattern of crossing over in Caenorhabditis elegans. Genetics. 1995;141(4):1339–1349.

[55] Chung G, Rose AM, Petalcorin MI, Martin JS, Kessler Z, Sanchez-Pulido L, et al. REC-1 and HIM-5 distribute meiotic crossovers and function redundantly in meiotic double-strand break formation in Caenorhabditis elegans. Genes & Development. 2015;29(18):1969–1979.

[56] Parée T, Noble L, Ferreira Goņcalves J, Teotónio H. rec-1 loss of function increases recombination in the central gene clusters at the expense of autosomal pairing centers. Genetics. 2023;p. iyad205.

[57] Teotónio H, Carvalho S, Manoel D, Roque M, Chelo IM. Evolution of out-crossing in experimental populations of Caenorhabditis elegans. PloS one. 2012;7(4):e35811.

[58] Chelo IM, Nédli J, Gordo I, Teotónio H. An experimental test on the probability of extinction of new genetic variants. Nature communications. 2013;4(1):2417.

[59] Theologidis I, Chelo IM, Goy C, Teotónio H. Reproductive assurance drives transitions to self-fertilization in experimental Caenorhabditis elegans. BMC biology. 2014;12:1–21.

[60] Noble LM, Chelo I, Guzella T, Afonso B, Riccardi DD, Ammerman P, et al. Polygenicity and epistasis underlie fitness-proximal traits in the Caenorhabditis elegans multiparental experimental evolution (CeMEE) panel. Genetics. 2017;207(4):1663–1685.

[61] Noble LM, Rockman MV, Teotónio H. Gene-level quantitative trait mapping in Caenorhabditis elegans. G3. 2021;11(2):jkaa061.

[62] Sved JA. Linkage disequilibrium and homozygosity of chromosome segments in finite populations. Theor Pop Biol. 1971;2:125–141.

[63] Lewontin RC. The interaction of selection and linkage. I. General considerations; heterotic models. Genetics. 1964;49(1):49.

[64] Charlesworth B. Directional selection and the evolution of sex and recombination. Genetics Research. 1993;61(3):205–224.

[65] Stetsenko R, Roze D. The evolution of recombination in self-fertilizing organisms. Genetics. 2022;222(1):iyac114.

[66] Proulx SR, Teotónio H. Selection on modifiers of genetic architecture under migration load. PLoS Genet. 2022;18(9):e1010350.

[67] Martin G, Otto SP, Lenormand T. Selection for recombination in structured populations. Genetics. 2006;172(1):593–609.

[68] Kamran-Disfani A, Agrawal AF. Selfing, adaptation and background selection in finite populations. J Evol Biol. 2014;27(7):1360–71.

[69] Johnston SE. Understanding the Genetic Basis of Variation in Meiotic Recombination: Past, Present, and Future. Molecular Biology and Evolution. 2024;41(7).

[70] Ubeda F, Russell TW, Jansen VAA. PRDM9 and the evolution of recombination hotspots. Theor Popul Biol. 2019;126:19–32.

[71] Li H, Lan R, Peng N, Sun J, Zhu Y. High resolution melting curve analysis with MATLAB-based program. Measurement. 2016;90:178–186.

[72] Farrar JS, Wittwer C. High-resolution melting curve analysis for molecular diagnostics. In: Molecular diagnostics. Elsevier; 2017. p. 79–102.

[73] R Core Team.: R: A Language and Environment for Statistical Computing. Vienna, Austria.

[74] Crow JF, Kimura M. An Introduction to Population Genetics Theory. New York: Harper Row, Publishers; 1970.

[75] Bates D, Maechler M, Bolker B, Walker S. Fitting Linear Mixed-Effects Models Using lme4. Journal of Statistical Software. 2015;67:1–48.

[76] Harris TW, Baran J, Bieri T, Cabunoc A, Chan J, Chen WJ, et al. WormBase 2014: new views of curated biology. Nucleic acids research. 2014;42(D1):D789–D793.

[77] Li H, Durbin R. Fast and accurate short read alignment with Burrows–Wheeler transform. bioinformatics. 2009;25(14):1754–1760.

[78] Danecek P, Bonfield JK, Liddle J, Marshall J, Ohan V, Pollard MO, et al. Twelve years of SAMtools and BCFtools. Gigascience. 2021;10(2):giab008.

[79] Garrison E, Marth G. Haplotype-based variant detection from short-read sequencing. arXiv preprint arXiv:12073907. 2012;.

[80] Meuwissen TH, Hayes BJ, Goddard ME. Prediction of total genetic value using genome-wide dense marker maps. Genetics. 2001;157(4):1819–29.

[81] Speed D, Hemani G, Johnson MR, Balding DJ. Improved heritability estimation from genome-wide SNPs. Am J Hum Genet. 2012;91(6):1011–21.

[82] Giovanny CP. Genome assisted prediction of quantitative traits using the R package sommer. PLoS ONE. 2016;11:1–15.

[83] Lenth RV. lsmeans: Least-Squares Means. R package version 2.20-23. http://CRANR-projectorg/package=lsmeans. 2015;.

[84] Hooper D. Handling, fixing, staining and mounting nematodes. 1986;.

[85] Stewart AD, Phillips PC. Selection and maintenance of androdioecy in Caenorhabditis elegans. Genetics. 2002;160(3):975–982.

[86] Swierczek NA, Giles AC, Rankin CH, Kerr RA. High-throughput behavioral analysis in C. elegans. Nature methods. 2011;8(7):592–598.

[87] Mallard F, Noble L, Guzella T, Afonso B, Baer CF, Teotónio H. Phenotypic stasis despite genetic divergence and differentiation in Caenorhabditis elegans. Peer Community in Evolutionary Biology. 2023;3:e119.

[88] Chen T, He T, Benesty M, Khotilovich V. Package ‘xgboost’. R version. 2019;90:1–66.

[89] Haller BC, Messer PW. SLiM 4: Multispecies Eco-Evolutionary Modeling. The American Naturalist. 2023;201(5):E127–E139.

[90] Lewontin R, Kojima Ki. The evolutionary dynamics of complex polymorphisms. Evolution. 1960;p. 458–472.

[91] Chelo IM, Nédli J, Gordo I, Teotónio H. An experimental test on the probability of extinction of new genetic variants. Nature communications. 2013;4(1):2417.

[92] Eisenmann DM, Kim SK. Protruding vulva mutants identify novel loci and Wnt signaling factors that function during Caenorhabditis elegans vulva development. Genetics. 2000;156(3):1097–1116.

[93] Hastie T, Tibshirani R. Exploring the nature of covariate effects in the proportional hazards model. Biometrics. 1990;p. 1005–1016.

